# Distal and proximal cis-regulatory elements sense X-chromosomal dosage and developmental state at the *Xist* locus

**DOI:** 10.1101/2021.03.29.437476

**Authors:** Rutger A.F. Gjaltema, Till Schwämmle, Pauline Kautz, Michael Robson, Robert Schöpflin, Liat Ravid Lustig, Lennart Brandenburg, Ilona Dunkel, Carolina Vechiatto, Evgenia Ntini, Verena Mutzel, Vera Schmiedel, Annalisa Marsico, Stefan Mundlos, Edda G. Schulz

## Abstract

Developmental genes such as *Xist*, the master regulator of X-chromosome inactivation (XCI), are controlled by complex *cis*-regulatory landscapes, which decode multiple signals to establish specific spatio-temporal expression patterns. *Xist* integrates information on X-chromosomal dosage and developmental stage to trigger XCI at the primed pluripotent state in females only. Through a pooled CRISPR interference screen in differentiating mouse embryonic stem cells, we identify functional enhancer elements of *Xist* during the onset of random XCI. By quantifying how enhancer activity is modulated by X-dosage and differentiation, we find that X-dosage controls the promoter-proximal region in a binary switch-like manner. By contrast, differentiation cues activate a series of distal elements and bring them into closer spatial proximity of the *Xist* promoter. The strongest distal element is part of an enhancer cluster ∼200 kb upstream of the *Xist* gene which is associated with a previously unannotated *Xist*-enhancing regulatory transcript, we named *Xert*. Developmental cues and X-dosage are thus decoded by distinct regulatory regions, which cooperate to ensure female-specific *Xist* upregulation at the correct developmental time. Our study is the first step to disentangle how multiple, functionally distinct regulatory regions interact to generate complex expression patterns in mammals.

## Introduction

During embryonic development, correct spatio-temporal gene expression is controlled by complex *cis*-regulatory landscapes (Bolt and Duboule, 2020). Multiple *trans*-acting signals in the form of sequence-specific transcription factors bind to *cis*-acting proximal and distal regulatory elements (RE) and control transcription from a gene’s core promoter, to precisely tune tissue- and stage-specific gene expression (Long et al., 2016; Spitz and Furlong, 2012). Another layer of regulation is composed of long non-coding RNAs (lncRNAs) that regulate neighboring genes in *cis* and are often transcribed from or through enhancer elements (Gil and Ulitsky, 2018). While *cis*-regulatory landscapes have been mapped for a number of genes (Fulco et al., 2016, 2019; Klann et al., 2017), it remains poorly understood how they decode complex information to precisely tune gene expression during development.

Here we use the murine *Xist* locus as a model to study information processing by *cis*-regulatory landscapes. *Xist* is an essential developmental regulator, which initiates X-chromosome inactivation (XCI) in females to ensure X-chromosome dosage compensation between the sexes (Brown et al., 1991; Penny et al., 1996). *Xist* is upregulated during early embryonic development from one out of two X chromosomes in females in an X-dosage dependent manner (Mutzel and Schulz, 2020). It then mediates chromosome-wide gene silencing in *cis* through successive heterochromatinization (Żylicz and Heard, 2020). The *Xist* locus must thus integrate differentiation cues and X-dosage information to establish the correct expression pattern.

In mice, XCI occurs in two waves. Shortly after fertilization the paternal X chromosome (Xp) is inactivated in an imprinted form of XCI, which is maintained in the extraembryonic tissues (Mak et al., 2004; Okamoto et al., 2004). In the inner cell mass of the blastocyst, which will give rise to the embryo, the Xp becomes reactivated again. Shortly after, at the primed pluripotent state, random XCI is initiated causing each cell to inactivate either the paternal or the maternal X. Random XCI, which is thought to occur in all placental mammals, can be recapitulated in cell culture by inducing differentiation of pluripotent mouse embryonic stem cells (mESCs) (Monk, 1981).

The regulatory landscape of *Xist*, called the X-inactivation center (*Xic*), is thought to encompass a region of ∼800 kb surrounding the *Xist* gene (Fig. 1a). The *Xic* is structured into two topologically associating domains (TADs), TAD-D and TAD-E, with the *Xist* gene being transcribed across their boundary (Nora et al., 2012). TAD-D contains several *Xist* repressors, including *Xist’s* non-coding antisense transcript *Tsix*, the *Tsix* enhancer region *Xite* and the more distal *Linx* locus (Galupa et al., 2020; Lee and Lu, 1999; Lee et al., 1999; Luikenhuis et al., 2001; Nora et al., 2012; Ogawa and Lee, 2003). TAD-E comprises the *Xist* promoter and multiple positive regulators, including two more lncRNA genes, *Jpx* and *Ftx*, which activate *Xist* expression and are upregulated concomitantly with *Xist* during mESC differentiation (Chureau et al., 2011; Furlan et al., 2018; Tian et al., 2010). In addition, TAD-E contains the protein-coding *Rnf12* gene, which contributes to X-dosage dependent *Xist* upregulation (Jonkers et al., 2009). While a series of *cis*- and *trans*-acting *Xist* activators have thus been identified, to our knowledge, no classical enhancer elements have been described to date.

**Figure 1.**
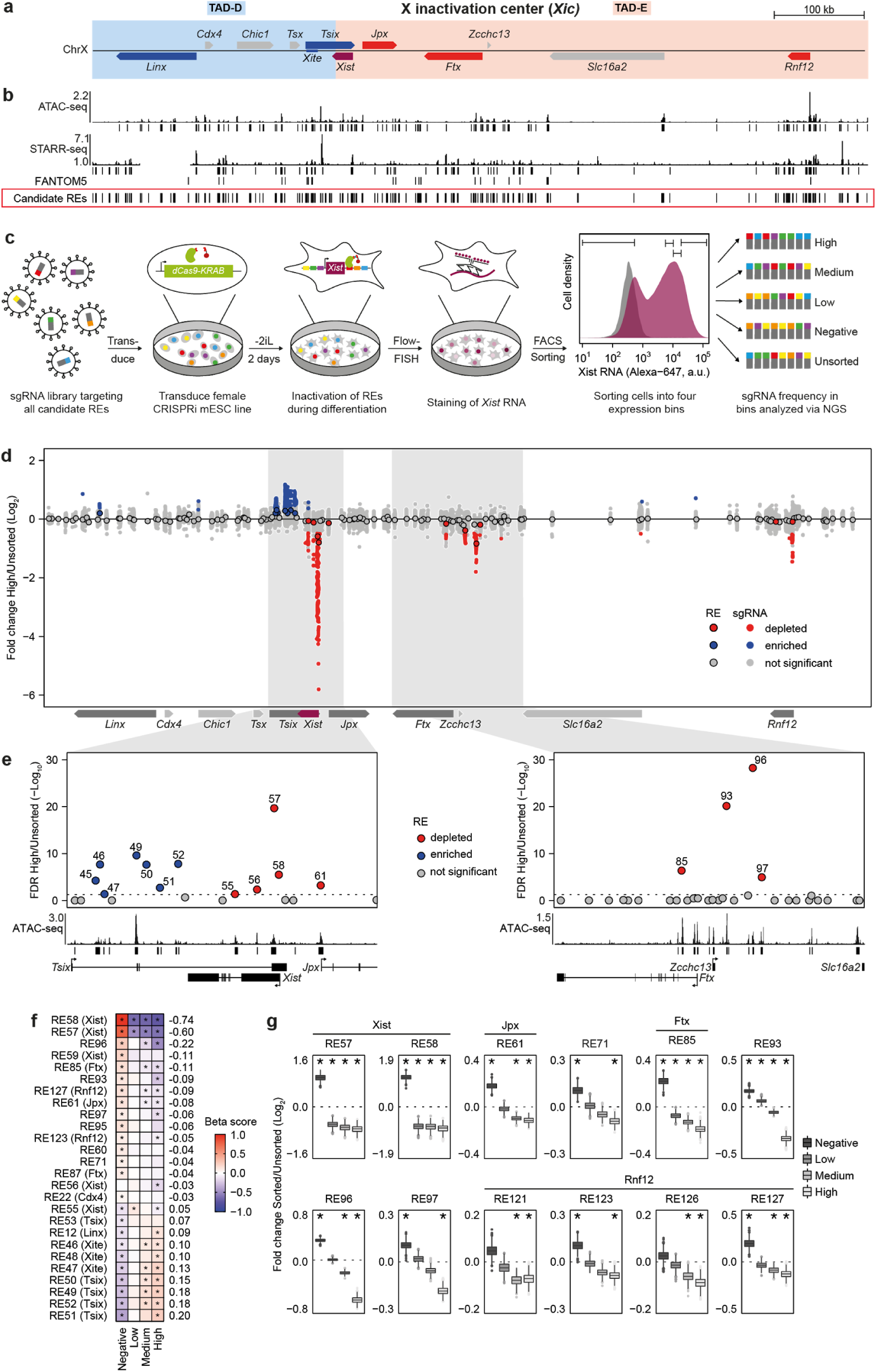
Identification of *Xist*-regulating genomic elements through a pooled CRISPR screen. (**a**) Schematic representation of the *Xic*, showing annotated transcripts and TAD positions (shaded boxes). Known *Xist* regulators in red (activators) and blue (repressors). (**b**) ATAC-seq, STARR-seq and FANTOM5 data used to identify candidate REs (red box). Vertical bars below the tracks represent peaks identified by MACS2 (FDR <0.1). A region within *Linx* was missing from the STARR-seq library. (**c**) Schematic outline of the pooled CRISPRi screen used to identify functional *Xist* REs in (d-g). After lentiviral transduction of female mESCs stably expressing an inducible dCas9-KRAB system were differentiated for two days by 2iL withdrawal and stained for *Xist* RNA by Flow-FISH (purple; undifferentiated cells grey). SgRNA distributions were analyzed in 4 expression bins each containing 15% of cells as indicated. (**d-e**) Comparison of sgRNA abundance in the Xist-high fraction compared to the unsorted population. Small dots in (d) show individual sgRNAs and rimmed circles in (d-e) denote results from a joint analysis of all sgRNAs targeting one RE. Significantly enriched and depleted sgRNAs (*MAGeCK test,* two-sided p-value <0.05) and REs (*MAGeCK mle,* Wald.FDR <0.05) are colored blue and red, respectively. The regions shaded in grey in (d) are shown in (e). In (**e**) significantly enriched or depleted REs are annotated with their number and ATAC-seq tracks from differentiated XX_ΔXic_ at day 2 are shown below the plot. The dashed line represents an FDR of 0.05. (**f**) Heatmap showing effect size estimated by *MAGeCK mle* (beta-score) when comparing each sorted fraction to the unsorted cells. All candidate REs significantly enriched or depleted in at least one sorted fraction (FDR < 0.05, asterisks) are shown. REs are sorted by their mean beta-score across all fractions (indicated on the right), with the score for the negative fraction negated. (**g**) Log2 fold-change of sorted and unsorted populations for 1000 bootstrap samples of 50 randomly selected sgRNAs for each RE. REs in TAD-E with an empirical FDR <0.01 (asterisks) in at least two populations are shown.

X-dosage information is in part transmitted through a double dose of RNF12 in female cells, which targets the *Xist* repressor REX1/ZFP42 for degradation (Gontan et al., 2012, 2018; Jonkers et al., 2009). REX1 is thought to repress *Xist* indirectly by enhancing *Tsix* transcription (Navarro et al., 2010), and directly through binding *Xist’s* transcription start site (TSS) and a CpG island ∼1.5 kb downstream of the TSS, where it competes for binding with the ubiquitous *Xist* activator YY1 (Chapman et al., 2014; Gontan et al., 2012; Makhlouf et al., 2014). Developmental regulation of *Xist* has been attributed to pluripotency factors (Donohoe et al., 2009; Navarro et al., 2008, 2010; Payer et al., 2013). They repress *Xist* in pluripotent cells, while their downregulation following differentiation triggers *Xist* upregulation. This pluripotency factor-induced repression is thought to be mediated by a pluripotency factor binding site within the first intron of *Xist,* together with transcriptional activation of *Tsix* (Donohoe et al., 2009; Navarro et al., 2008, 2010). However, neither deletion of the intronic binding site nor of the *Tsix* promoter results in *Xist* upregulation prior to differentiation (Barakat et al., 2011; Lee and Lu, 1999; Minkovsky et al., 2013; Nesterova et al., 2011). It thus remains an open question how the developmental state is sensed by the *Xist* locus and whether developmental regulation is indeed ensured through pluripotency factor repression alone or whether differentiation cues might also activate *Xist*.

To understand how the complex *cis*-regulatory landscape of *Xist* integrates information on X-dosage and development, we have comprehensively mapped *cis*-regulatory elements that control *Xist* in mESCs. We then profiled how their activity is modulated by X-dosage and differentiation. In this way, we identified an enhancer cluster that controls developmental *Xist* upregulation and is associated with a previously unannotated transcript we named *Xert*. We show that *Xert* is a lncRNA that activates *Xist* transcription in *cis* and that the locus interacts with the *Xist* promoter in 3D-space. Overall, our data show that differentiation cues are integrated by a series of distal regulatory elements. They can however stimulate *Xist* transcription only in female cells, where double X-dosage acts to prevent repression of the promoter-proximal region.

## Results

### Identification of *cis*-regulatory elements that control *Xist* through a pooled CRISPR screen

To understand how information is processed by *Xist*’s regulatory landscape, we comprehensively identified REs that control *Xist* upregulation at the onset of random XCI. We performed a pooled CRISPR interference (CRISPRi) screen (Klein et al., 2018), where catalytically dead Cas9 (dCas9) fused to a KRAB repressor domain is targeted to putative REs in a pooled fashion to inactivate one RE per cell. Subsequent enrichment of cells with high or low *Xist* expression allows identification of functional REs by comparing sgRNA abundance between cell populations.

To establish a set of candidate REs to be tested in the screen, we profiled DNA accessibility and enhancer activity in an episomal reporter assay within the *Xic*. To this end, we performed ATAC-seq and STARR-seq in naive and differentiating conditions (Fig. 1b, Supplementary Fig. 1a, see methods for details) (Arnold et al., 2013; Buenrostro et al., 2013). After integrating these data sets with enhancer regions reported by the FANTOM5 consortium (Lizio et al., 2015), we defined a total of 138 candidate REs with a median length of 991 bp (Fig. 1b, Supplementary Fig. 1b-c). SgRNAs targeting those candidate REs were cloned into a lentiviral vector, resulting in a library with 7358 sgRNAs and a median number of 43 guides per RE (Supplementary Fig. 1d-e, Supplementary Table 1).

A female mESC line (TX-SP107) stably expressing an abscisic acid (ABA) inducible CRISPRi system (Gao et al., 2016) was transduced with the sgRNA library, differentiated for 2 days by 2i/LIF (2iL) withdrawal and stained for *Xist* RNA by Flow-FISH (Fig. 1c, Supplementary Fig. 1f-g). Cells were sorted into 4 populations (negative, low, medium, high) according to their *Xist* levels (Fig. 1c). The relative sgRNA frequency within the unsorted and sorted populations was determined by deep sequencing (Supplementary Fig. 1h-j).

To identify REs controlling *Xist*, we compared sgRNA abundance between sorted and unsorted populations using the *MAGeCK* tool suite (Fig. 1d-e, Supplementary Fig. 1k-l, Supplementary Table 2) (Li et al., 2014, 2015). All regions within the *Xic* that had previously been described to activate *Xist*, were depleted from the *Xist*-high fraction and enriched in the negative population, while known repressive elements showed the opposite pattern (see Supplementary Screen discussion). Among others the screen identified the *Xist* promoter (RE58), the promoter-proximal CpG island (RE57) (Johnston et al., 1998), the *Jpx* promoter (RE61) (Tian et al., 2010), the *Ftx* promoter region (RE85, RE87) (Chureau et al., 2011; Furlan et al., 2018) and multiple regions within *Tsix* (RE46-50) (Lee and Lu, 1999; Ogawa and Lee, 2003) (Fig. 1d-e, Supplementary Fig. 1k-l). Importantly, these regions do not necessarily represent *Xist* enhancer elements. Since most of them contain the TSS of known *Xist* regulators, the observed effects might rather be mediated by repression of the linked transcript.

In addition to these known elements, the screen also identified several regions that, to our knowledge, have not yet been shown to regulate *Xist*. Multiple intronic elements within *Tsix* (RE51-53) had repressive effects, and an element downstream of *Rnf12* (RE123), which might act as an *Rnf1*2 enhancer, activated *Xist* expression (Fig. 1d-e, Supplementary Fig. 1k-l). The most prominent region identified through our screen was a cluster of activating REs (RE93, 95-97) ∼150-170 kb telomeric of *Xist*, which were all enriched in the *Xist*-negative population and all except RE95 were also depleted from *Xist*-high cells.

We next ranked all REs according to their contribution to *Xist* regulation using two different approaches (Fig. 1f, Supplementary Table 2, see methods for details). As expected, the strongest activating regions were located around the *Xist* promoter, most notably at the TSS (RE58) and the promoter-proximal CpG island (RE57). Among the distal elements, the newly discovered RE96 region showed the strongest effect, followed by a region in *Ftx* (RE85) and another previously unknown element RE93 (Fig. 1f). Interestingly, we observed distinct enrichment patterns among elements, across the different Xist-positive populations. While promoter-proximal REs (RE57, 58) were depleted to a similar extent across populations, most distal elements, in particular the newly identified RE93-97 region, showed a gradual increase in depletion from the Xist-low to high populations (Fig. 1g, Supplementary Fig. 1m). This suggests that the promoter-proximal elements control *Xist* in a binary fashion, thus constituting an ON/OFF switch. By contrast, the distal RE93-97 elements seem to control expression levels once the *Xist* promoter has been switched on.

### Proximal and distal elements integrate X-dosage information and differentiation cues

In the next step, we investigated how activity of the identified *Xist*-controlling REs was modulated by differentiation and X-chromosomal dosage. To this end, we profiled a series of histone modifications and DNA accessibility before and during differentiation in mESCs with one (XO) and two X chromosomes (XX), but otherwise identical genetic backgrounds (Fig. 2a). To unequivocally distinguish the inactive X (Xi) and the active X (Xa), we used a female mESC line (XX_ΔXic_) with a monoallelic ∼800 kb deletion around *Xist* (Fig. 2a) (Pacini et al., 2020). Only the *Rnf12* gene at the distal end of TAD-E remained intact to not preclude *Xist* upregulation from the wildtype allele (Barakat et al., 2014). *Xist* expression was only slightly reduced in XX_ΔXic_ cells compared to the parental TX1072 line, with >70% of cells expressing *Xist* at day 2-4 of differentiation (Supplementary Fig. 2a-b). We profiled 7 histone modifications (H3K4me3, H3K27ac, H3K4me1, H3K9me3, H3K27me3, H2AK119ub, H3K36me3) through CUT&Tag (Kaya-Okur et al., 2019), under naive conditions and at day 2 (when Xist is strongly upregulated) and at day 4 of differentiation (when gene silencing is established) (Pacini et al., 2020). The data showed the expected peak patterns and was in good agreement with native ChIP-seq in the parental line (Supplementary Fig. 2c-g, Supplementary Table 3) (Żylicz et al., 2019).

**Figure 2.**
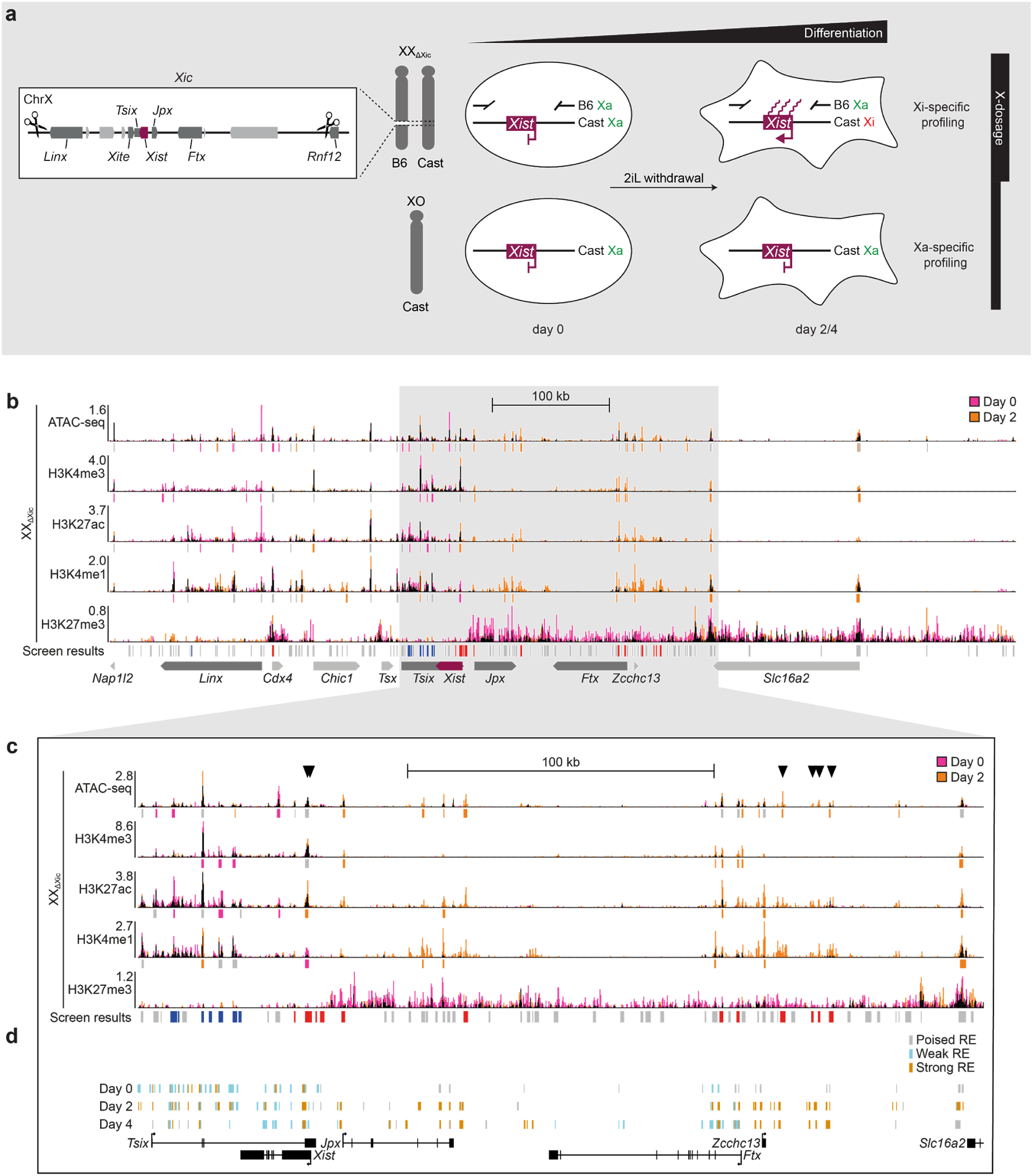
Differentiation cues activate distal, but not proximal *Xist*-controlling elements. (**a**) Schematic representation of the experimental system to profile the Xi and Xa in XX and XO mESCs, respectively. Deletion of the *Xic* (chrX: 103182701-103955531) from one X chromosome in female XX_ΔXic_ (TXΔXic_B6_) mESCs (left) allows chromatin profiling of the *Xist*-expressing chromosome upon differentiation by 2iL withdrawal in comparison to an XO line, where *Xis*t will stay silent (right). (**b-c**) DNA accessibility and histone modifications in female XX_ΔXic_ mESCs (a) prior to (Day 0) and during differentiation (Day 2) profiled by ATAC-seq and CUT&Tag. In the track overlay signal specific for day 0 and day 2 is colored in pink and orange as indicated. Reads from two biological replicates were merged. Vertical bars below the tracks (except for H3K27me) mark peaks called in at least one time point and are colored (pink/orange), if the signal is significantly different (FDR <0.05) between the time points across both biological replicates. The screen results (Fig. 1) are shown below the tracks, where candidate REs that inhibit (blue) or activate (red) *Xist* expression in the negative or high fractions of the CRISPR screen are colored. The entire deleted region (b) and a zoom-in (c) of Xist and its upstream region (indicated in grey in b) are shown. Arrowheads in (c) indicate the promoter-proximal elements and the distal enhancers RE93,95-97. (**d**) Chromatin segmentation using *ChromHMM* based on the data in (c). Only regions classified as REs are shown and colored as indicated.

Although *Xist* was strongly upregulated between day 0 and day 2 in XX_ΔXic_ cells (Supplementary Fig. 2c), few changes occurred at its promoter-proximal region. It was devoid of repressive marks already in naive cells, exhibited DNA accessibility and was decorated by active histone modifications, such as H3K4me3, H3K4me1 and H3K27ac (Fig. 2b-c). Only a small but significant increase in H3K27ac and loss of the H3K4me1 mark was observed together with a reduction of H3K36me3 (Fig. 2c, Supplementary Fig. 2i). The latter likely reflects *Tsi*x downregulation, which is thought to repress *Xist* by co-transcriptional deposition of this mark (Loos et al., 2015; Ohhata et al., 2015). The *Xist* promoter thus resides in a “poised” state already prior to differentiation. In contrast to promoter-proximal elements, the distal REs we had identified in the screen (Ftx, RE93-97) were largely inactive under naive conditions and gained active chromatin marks and DNA accessibility only during differentiation (Fig. 2b-c). This observation was confirmed by chromatin segmentation with ChromHMM (Fig. 2d, Supplementary Fig. 2h) (Ernst and Kellis, 2012). Moreover, the distal elements were covered by a broad H3K27me3 domain in naive cells (Fig. 2b-c, Supplementary Fig. 2j), which corresponds to a previously described “H3K27me3 hotspot” (Marks et al., 2009; Rougeulle et al., 2004). As previously reported, the hotspot disappeared during differentiation (Marks et al., 2009), potentially contributing to the observed activation of the Ftx-RE93-97 region (Fig. 2b-c, Supplementary Fig. 2j). These results suggest that *Xist* upregulation during differentiation is primarily driven by distal regulatory elements.

When comparing distal REs between XX_ΔXic_ and XO cells we found that they gained active marks and lost H3K27me3 in a similar manner in both cell lines, suggesting that they are not controlled by X-chromosomal dosage (Fig. 3a-c, Supplementary Fig. 3a-c). The only regions that showed higher activity in XX_ΔXic_ than in XO cells at day 2 were the *Xist* promoter-proximal elements (Fig. 3a-d). While they appeared mostly active in both cell lines at day 0, they lost activity in XO cells during differentiation (Fig. 3b-d). Concomitantly, a broad ∼16 kb-wide H3K9me3 domain, covering the *Xist* promoter region, appeared only in XO cells at day 2 and 4 (Fig. 3b, Supplementary Fig. 3c). X-chromosomal dosage thus appears to mainly control the promoter-proximal region, where it counteracts active repression by H3K9me3 during differentiation. Developmental cues, on the other hand are primarily sensed by distal regulatory elements.

**Figure 3.**
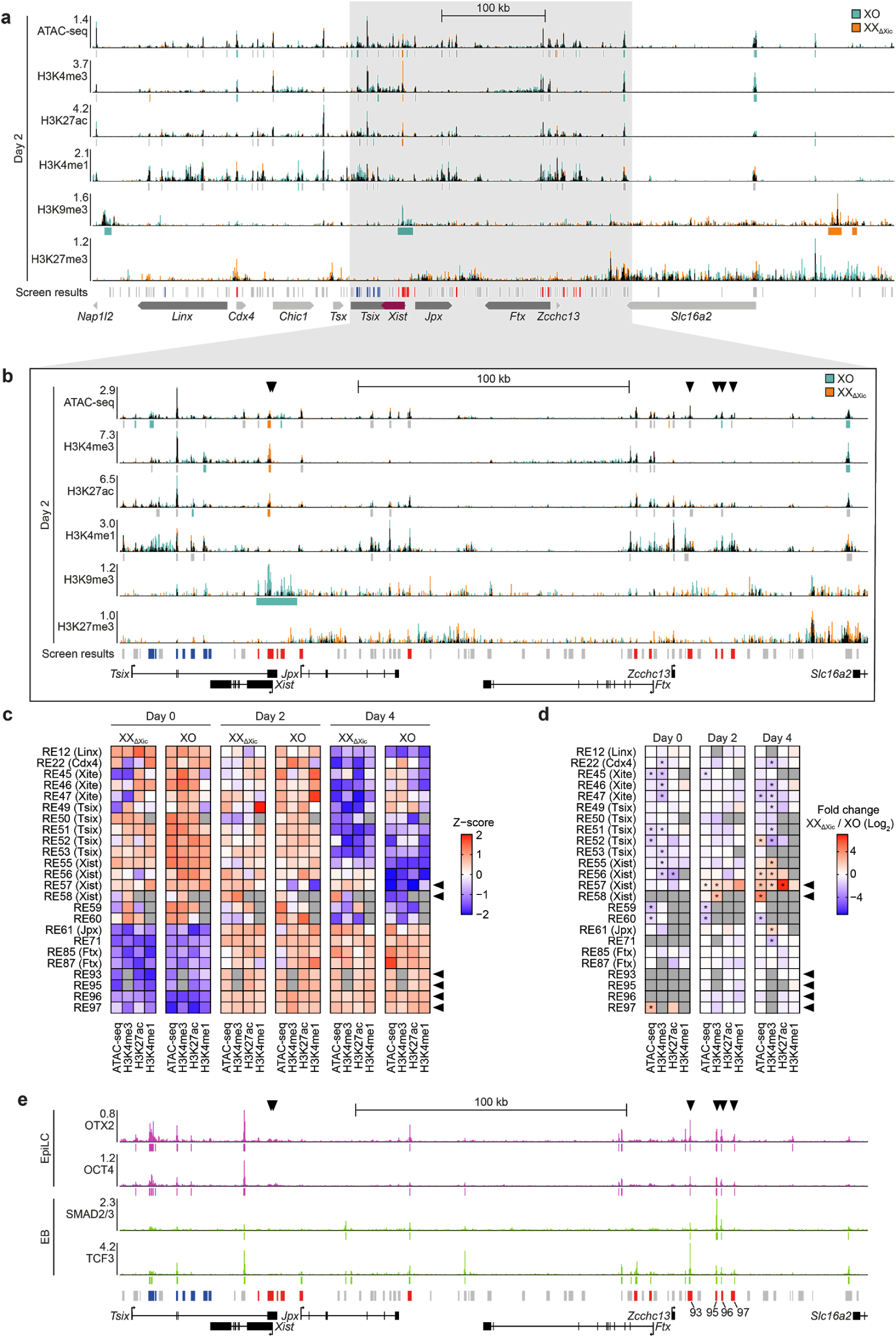
X-dosage information is decoded by promoter-proximal elements. (**a-b**) DNA accessibility and histone modifications in XX_ΔXic_ and XO mESCs after 2 days of differentiation as in Fig. 2b-c. Positions with increased signal in XX_ΔXic_ or XO mESCs are indicated in orange and teal, respectively. (**c-d**) Quantification of the indicated data sets within REs found to regulate *Xist* in the CRISPR screen. REs with insufficient coverage are grayed out. In (d) significant differences (p <0.05) according to *DiffBind* analysis are marked with an asterisk. (**e**) Published ChIP-seq tracks (Buecker et al., 2014; Wang et al., 2017) for OTX2, OCT4, SMAD2/3 and TCF3 in epiblast-like cells (EpiLCs) or embryoid bodies (EBs). Arrowheads in (b-e) indicate the promoter-proximal elements and the distal enhancers RE93,95-97.

To further investigate what drives activation of distal REs, in particular RE93-97, we aimed to identify transcription factors that might regulate these enhancer elements. We used the Cistrome database, which contains a large collection of published ChIP-seq experiments in different cell types and tissues (Zheng et al., 2019), to identify factors that were enriched at RE93, 95, 96 and 97 (Supplementary Fig. 3d). All 4 REs were bound by OTX2, a factor that regulates the transition between the naive and primed pluripotent states (Fig. 3e) (Acampora et al., 2013; Yang et al., 2014). Mechanistically, OTX2 induces repositioning of the pluripotency factor OCT4 (Buecker et al., 2014), which also binds the RE93-97 region in epiblast-like cells (EpiLC), but not in mESCs (Fig. 3e, Supplementary Fig. 3e). Moreover, we detected binding of two other regulators of ESC differentiation, SMAD2/3 and TCF3 (Guo et al., 2011; Pauklin and Vallier, 2015), specifically in differentiated cells (embryoid bodies, Fig. 3e, Supplementary Fig. 3e) (Wang et al., 2017).

In sum, the *Xist* promoter is already in a mostly active chromatin configuration prior to differentiation, while distal enhancers are not yet active and are covered by a broad repressive H3K27me3 domain. These distal elements are then activated by several differentiation-associated transcription factors in XX_ΔXic_ and XO cells, but *Xist* upregulation appears to be prevented in XO cells through H3K9me3 deposition at the *Xist* promoter.

### A long non-coding RNA named *Xert* is transcribed through the distal enhancer cluster concomitantly with Xist upregulation

To investigate transcriptional activity at the newly identified distal enhancer regions, we profiled nascent transcription and mature RNA expression in our XX_ΔXic_-XO model. To this end, we performed transient transcriptome sequencing (TT-seq, Schwalb et al., 2016) and RNA-sequencing in naive cells and after 2 and 4 days of differentiation (Fig. 4a-b, Supplementary Fig. 4a-b, Supplementary Table 4). We detected an unannotated transcript, which overlapped with the *Xist* enhancers RE93-97 and was expressed upon differentiation in both cell lines (Fig. 4a, grey box). Through polyA-enriched RNA-seq as well as 3’- and 5’RACE we identified several relatively short (400-800 bp), spliced and poly-adenylated transcripts originating from a ∼50 kb genomic region (Fig. 4c, Supplementary Fig. 4c-d). They showed limited protein-coding potential, supporting a classification as lncRNAs (Supplementary Table 5). The main TSS was located within the RE93 element and exhibited a chromatin state typical for enhancers. It was characterized by chromatin accessibility, bidirectional transcription, H3K27ac and a high H3K4me1-to-H3K4me3 ratio (Fig. 4c-d), which is reminiscent of a previously described lncRNA class that are transcribed from enhancer elements (Gil and Ulitsky, 2018; Marques et al., 2013; Tan et al., 2020). Since the promoter of this unknown transcription unit was identified as an *Xist* enhancer element in our screen, we hypothesized that it might constitute a lncRNA that activates *Xist* transcription, similar to *Jpx* and *Ftx*. We thus named the locus *Xist-enhancing regulatory transcript* (*Xert*).

**Figure 4.**
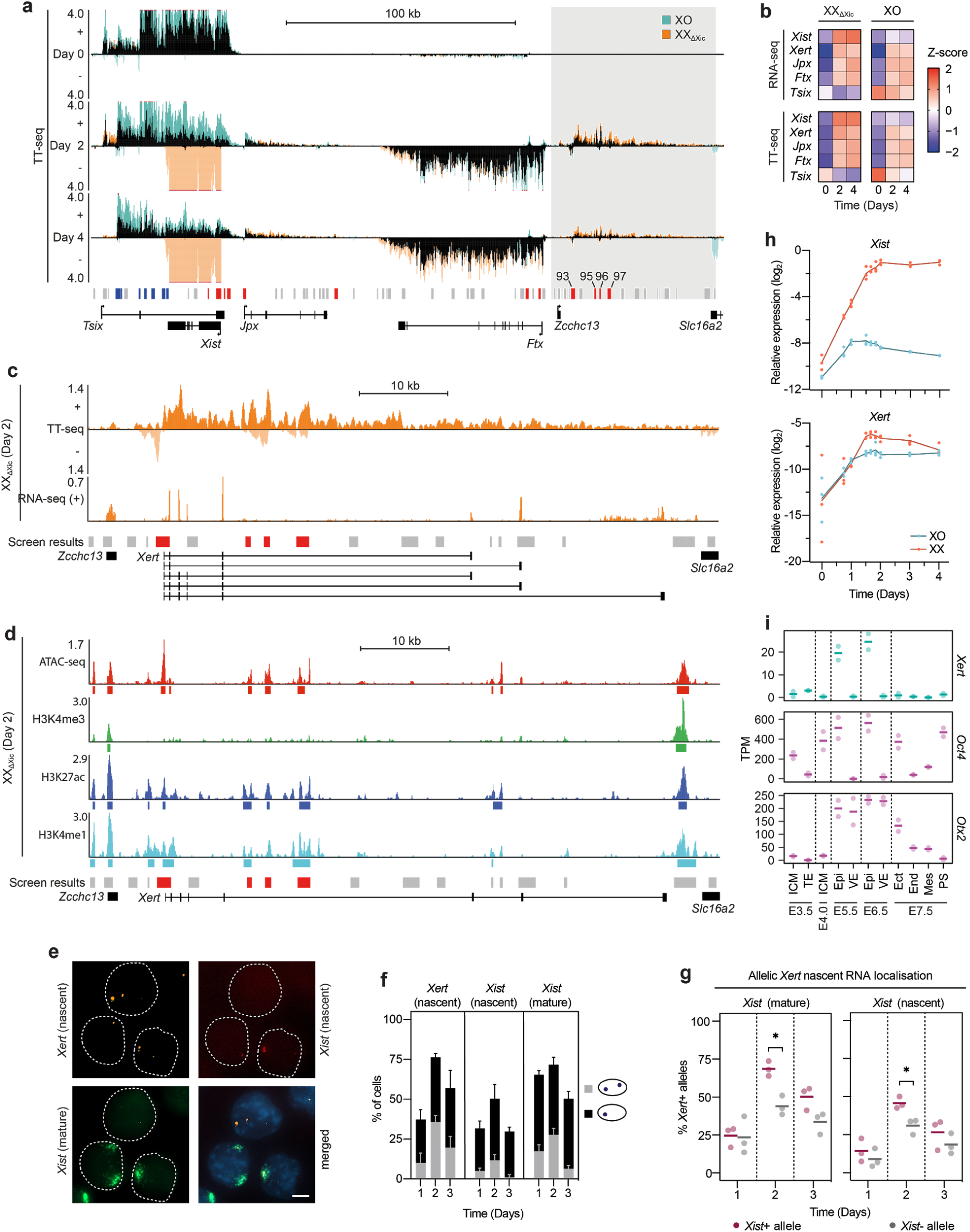
An unannotated enhancer-associated transcript is upregulated concomitantly with *Xist* at the onset of XCI. (**a-b**) TT-seq (a,b) and RNA-seq (b) in XX_ΔXic_ and XO cells before (Day 0) and during differentiation (Day 2, Day4). TT-seq coverage track in (a) shows nascent RNA transcription on the + and - strand; positions where the signal extends beyond the depicted range are marked in red. Grey box indicates an unannotated transcript and is enlarged in (c). The screen results (Fig. 1) are shown below the tracks, where candidate REs that inhibit (blue) or activate (red) *Xist* expression in the negative or high fractions of the CRISPR screen are colored. (**c**) TT-seq (both stands) and pA-RNA-seq (+ strand only) tracks of the region shaded in grey in (a) with five isoforms of the newly identified *Xert* transcripts (bottom). (**d**) DNA accessibility and histone modifications in differentiating XX_ΔXic_ cells at the *Xert* locus. (**e-g**) RNA-FISH in differentiating TX1072 cells. An example image (e) is shown for day 2, where nuclei are denoted by a dashed outline and scale bar marks 5 μm. Quantification of monoallelic (1 signal) and biallelic (2 signals) expression (f) and of the frequency of *Xert* detection at *Xist*- or *Xist*+ alleles in three biological replicates (g). Asterisks indicate significance of *p*<0.05 using an unpaired two-tailed *t*-test. (**h**) Expression kinetics of *Xist* and *Xert* in XX and XO cells during differentiation, measured by RT-qPCR (n=3). The line connects the mean, dots represent individual replicates. (**i**) *Xert, Otx2 and Oct4 (Pou5f1)* RNA expression during early mouse development (E3.5-E7.5) from embryos of both sexes combined (Zhang et al., 2018). Inner cell mass (ICM), trophectoderm (TE), epiblast (Epi), visceral endoderm (VE), ectoderm (Ect), endoderm (End), mesoderm (Mes), primitive streak (PS). In (g) and (i) the horizontal bar indicates the mean and dots individual replicates. In (f) mean and s.d. of 3 biological replicates are shown. 100 cells were quantified in each sample.

To further characterize *Xert*, we assessed its expression dynamics both *in vitro* and *in vivo*. First, we performed RNA-FISH for *Xert* and *Xist* in differentiating female mESCs (Fig. 4e-f). We found that *Xert* was more frequently detected on *Xist*-positive than on Xist-negative alleles, suggesting that it might activate *Xist* in *cis* (Fig. 4g). Following this, we performed a high-resolution time course experiment, which revealed that *Xert* was upregulated concomitantly with *Xist*, *Jpx* and *Ftx* at the onset of differentiation (Fig. 4h, Supplementary Fig. 4f). *Xert* reached ∼4-fold higher levels in XX compared to XO cells (and 1.5-1.8 times higher levels in XX_ΔXic_ compared to XO, Supplementary Fig. 4b). Therefore not only differentiation cues, but also X-chromosomal dosage appeared to modulate its expression levels. In contrast to *Ftx* and *Jpx*, which maintain high expression throughout the time course, *Xert* levels started to decrease after day 2, suggesting a role in initial *Xist* upregulation (Fig. 4h, Supplementary Fig. 4e). We next reanalyzed published data sets to characterize activity of the *Xert* region in mouse embryos. RNA-seq data from sex-mixed embryos (Yang et al., 2019) revealed that *Xert* was specifically expressed at the onset of random XCI, which occurs in the epiblast at embryonic days 5.5 and 6.5 (Fig. 4i, Supplementary Fig. 4f) (Mak et al., 2004; Shiura and Abe, 2019). The *Xert* expression pattern mirrored co-expression of *Xert* binding factors *Otx2* and *Oct4* (Fig. 3e, Fig. 4i), further supporting a role of these factors in regulating *Xert* expression. Re-analysis of ChIP-seq data from post-implantation embryos (Zhang et al., 2018) showed that the *Xert* promoter and the RE95-97 enhancer region located in its longest intron were marked with an active enhancer signature (H3K4me1 and H3K27ac) in the E6.5 epiblast with levels decreasing at E7.5 (Supplementary Fig. 4g).

Taken together, we have identified a lncRNA within the *Xic*, which is associated with a series of functional *Xist*-activating elements. Since it is specifically expressed at the onset of random XCI and is positively correlated with *Xist* transcription, it might function as an early *cis*-acting *Xist* activator.

### *Xert* transcription activates *Xist* in *cis*

To test a functional role of *Xert* in *Xist* regulation, we perturbed *Xert* transcription through multiple approaches (Fig. 5a). First, we attenuated *Xert* promoter (XertP) activity in female cells using CRISPRi (Fig. 5b-c). We observed a ∼20-fold reduction of *Xert* levels, resulting in ∼2-fold reduced *Xist* expression at day 2 (Fig. 5b, right). Flow-FISH revealed a 10-20% decrease in *Xist*-expressing cells with a 25% reduction of *Xist* levels within the positive population (Fig. 5c). Next, we overexpressed *Xert* in male cells using the SunTag CRISPR activation (CRISPRa) system (Heurtier et al., 2019; Tanenbaum et al., 2014) in a mESC line carrying a *Tsix* mutation to facilitate ectopic *Xist* upregulation (Supplementary Fig. 5a-b). Targeting the XertP region with three guides led to a ∼2-fold over-expression of *Xert* at day 2 of differentiation, resulting in a similar increase in *Xist* levels (Fig. 5d). We thus concluded that the *Xert* promoter region promotes *Xist* expression in differentiating mESCs.

**Figure 5.**
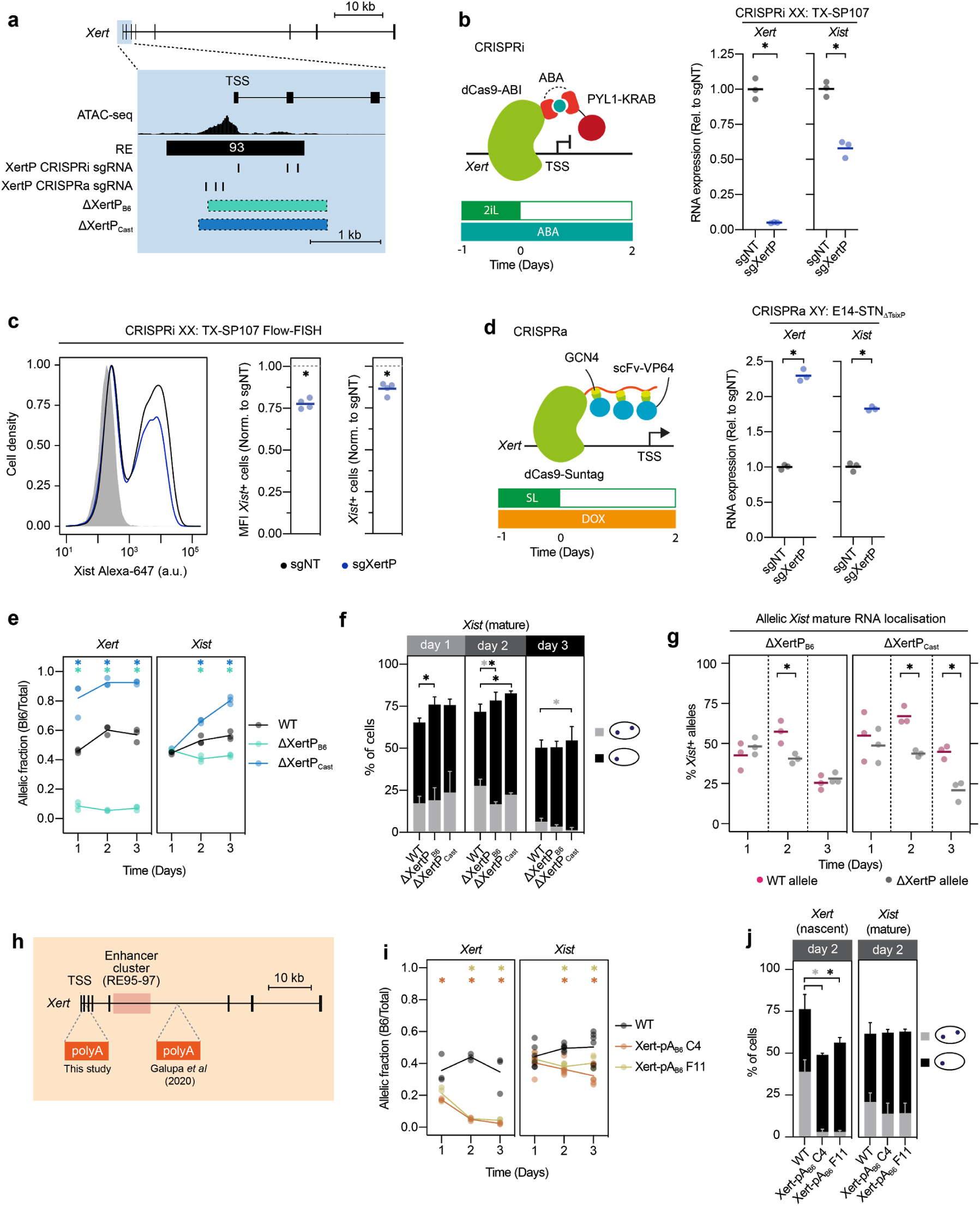
*Xert* transcription enhances *Xist* expression in *cis.* (**a**) Schematic of *Xert* promoter (XertP) perturbations employed in (b-g). (**b-d**) Repression of XertP through an ABA-inducible CRISPRi system in female TX-SP107 mESCs (b-c) and XertP activation through a Dox-inducible CRISPRa system in male E14-STN_ΔTsixP_ cells (d). Cells were transduced with multiguide expression vectors of three sgRNAs against XertP as indicated in (a) or with non-targeting controls (NT). RT-qPCR (b,d) and Flow-FISH (c) of 3-4 biological replicates after 2 days of differentiation are shown. Values of sgNT were calculated as the geometric mean of 4 (b,d) or 2 (c) different multiguide NT plasmids. In (c) the sample shaded in grey denotes undifferentiated (2iL) TX-SP107 cells. (**e-g**) Differentiation time course of two heterozygous XertP deletion lines (position shown in a) and the parental wildtype line (WT) assessed by pyrosequencing (e) and RNA-FISH (f-g). (f) Quantification of monoallelic (grey) and biallelic (black) expression of mature *Xist* RNA. 100 cells were quantified per replicate. (g) Frequency of *Xist* upregulation from the wildtype or the deleted allele identified through the presence or absence of an *Xert* signal, respectively. Only cells with a single *Xert* signal (14-66 cells per replicate) were included in the analysis. (**h-j**) Differentiation time course of two cell lines (Xert-pA_B6_) with premature termination of the *Xert* transcript through heterozygous insertion of a polyA signal (h) and the parental wildtype line (WT), assessed by pyrosequencing (i) and RNA-FISH (j). Horizontal bars (b-d,f) or lines (g,i) denote the mean of 3 (b,d,f,g), 4 (c) or 6 (i, *Xist*) biological replicates, dots represent individual measurements. In (e) and (j) mean and s.d. of 3 biological replicates are shown. Asterisks indicate significance of *p*<0.05 using an unpaired two-tailed *t*-test, or for (c) of a one-sample two-tailed *t*-test. Colored asterisks in (g) and (i) denote comparison of the respective mutant with the wildtype control.

To test whether *Xert* regulates *Xist* in *cis* or in *trans* we deleted the XertP region on one allele in female mESCs and assessed the effect on *Xist* expression. We generated two cell lines with a heterozygous deletion either on the Cast or B6 allele in female mESCs (ΔXertP), encompassing the accessible region upstream of the *Xert* TSS and the first two exons (Fig. 5a, Supplementary Fig. 5c-f). Monoallelic transcription of *Xert* was confirmed by RNA-FISH and by pyrosequencing, which performs quantitative sequencing over single SNPs on cDNA (Fig. 5e, Supplementary Fig. 5g-h). For *Xist*, the deletion led to a slight reduction of overall expression levels, accompanied by a shift from biallelic to monoallelic expression, with *Xist* being preferentially detected at the wildtype X chromosome (Fig. 5f-g, Supplementary Fig. 5h). Similarly, pyrosequencing revealed that 65-80% of *Xist* RNA in ΔXertP cells originated from the wildtype allele compared to 50% in the parental cell line (Fig. 5e). These results show that the *XertP* region enhances *Xist* transcription in *cis*.

Next, we aimed to distinguish whether the effects of perturbing the XertP element could be attributed to a regulatory role of *Xert* transcription or to the DNA element containing its promoter. We thus terminated transcription ∼1.2 kb downstream of the TSS through insertion of a polyadenylation (pA) cassette in one allele in female mESCs (Fig. 5h, Supplementary Fig. 5i-k). RNA-FISH and pyrosequencing confirmed efficient transcription termination (Fig. 5i-j, Supplementary Fig. 5l). Pyrosequencing revealed that *Xist* expression was skewed towards the wildtype allele (Fig. 5i) to a similar extent as in ΔXertP_B6_ mutant mESCs, where the same allele is modified (Fig. 5e). This finding suggests that *Xert* transcription plays a functional role in *Xist* regulation.

In summary, we could show that *Xert* is a *cis*-acting *Xist* activator, since promoter deletion and premature termination of the transcript resulted in reduced *Xist* upregulation from the mutated allele. Of note, in a previous study unrelated to *Xert* transcription, a pA signal was inserted at a more downstream location within *Xert* (Fig. 5h) without detectable effect on *Xist* (Galupa et al., 2020). This suggests that transcription through the RE95-97 enhancer cluster, which is located between the two insertion sites (Fig. 5h), might mediate *Xert*-induced *Xist* activation. We thus went on to characterize the functional role of this enhancer cluster in *Xist* regulation.

### *Xert*-associated enhancer elements control *Xist* upregulation

RE95-97 were identified as *Xist*-activating regions in our screen with RE96 exhibiting the strongest effect out of all distal elements in the *Xic* (Fig. 1f). We termed this region *Xert*-associated enhancer cluster (XertE). To confirm a functional role of XertE in *Xist* upregulation, we targeted each RE within the cluster individually with CRISPRi, which resulted in up to 3-fold reduced *Xist* expression after two days of differentiation (Fig. 6a-c).

**Figure 6.**
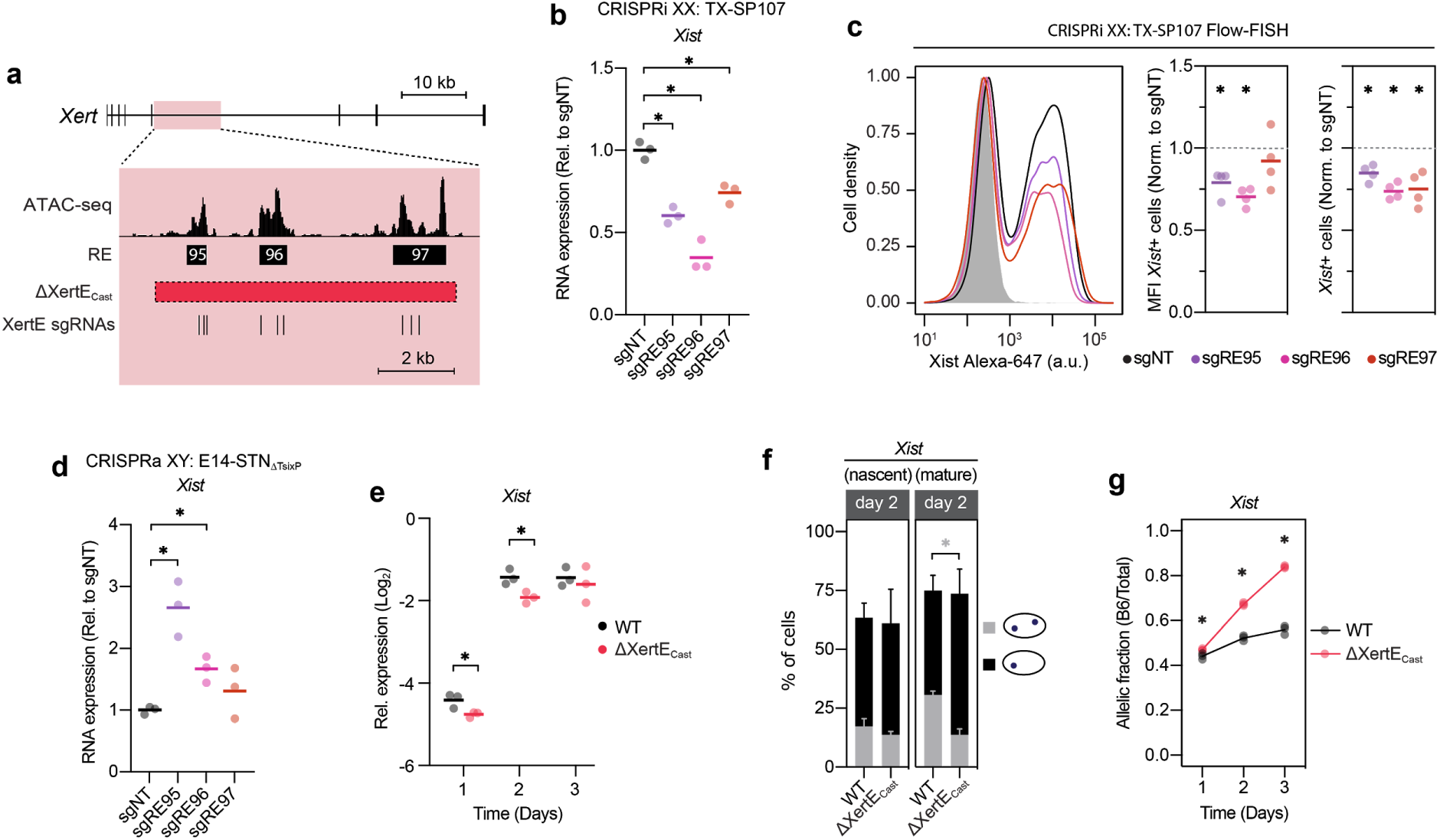
An intronic enhancer cluster within *Xert* activates *Xist* expression in *cis*. (**a**) Schematic of *Xert* enhancer cluster (XertE) perturbations used in (b-g). (**b-c**) Inducible CRISPRi repression of individual REs in XertE in differentiated (Day 2) TX-SP107 cells with stable multiguide expression of three sgRNA against each RE (position shown in a) or non-targeting controls (NT). *Xist* expression is quantified by RT-qPCR (b) and Flow-FISH (c) and normalized to sgNT, where each sgNT replicate is given by the geometric mean of four different sgNT plasmids. In (c) the sample shaded in grey denotes undifferentiated (2iL) TX-SP107 cells. (**d**) Ectopic activation of individual REs in XertE through an inducible CRISPRa Suntag system in male E14-STN_ΔTsixP_ cells with stable multiguide expression of three sgRNAs against each RE (positions shown in a). *Xist* expression was quantified after 2 days of differentiation by RT-qPCR and normalized to non-targeting controls (sgNT), where each sgNT replicate is given by the geometric mean of four different sgNT plasmids. (**e-g**) Characterization of a female mESC line, carrying a deletion of the XertE region (ΔXertE_Cast_) on the Cast X chromosome (position shown in a) and an inversion on the other allele. *Xist* expression was quantified by RT-qPCR (e), RNA-FISH (f) and pyrosequencing (g) in comparison to the parental TX1072 cell line (WT). Horizontal bars (b-e) or lines (g) denote the mean of 3 (b,d,e,g) or 4 (c) biological replicates, dots represent individual measurements. In (f) mean and s.d. of 3 biological replicates are shown. Asterisks indicate significance of *p*<0.05 using an unpaired two-tailed *t*-test, or for (c) of a one-sample two-tailed *t*-test.

In addition,we activated the XertE region with the same sgRNAs through CRISPRa, which has previously been used to interrogate enhancer function (Li et al., 2020). Activation of the XertE region resulted in increased *Xist* expression in male and female cells (Fig. 6d, Supplementary Fig. 6a), confirming a functional role of XertE in *Xist* upregulation. To further characterize XertE we deleted an ∼8 kb region containing all three elements. Due to repetitive sequences around the target region it was difficult to identify correctly modified clones. After several optimization attempts, we identified one clone that carried the deletion on the Cast allele (ΔXertE_Cast_), but also an inversion of the region on the B6 chromosome (Fig. 6a, Supplementary Fig. 6b-d). Since enhancers are thought to function in an orientation-independent manner, we expected the inverted allele to behave like a wildtype and went on to characterize this cell line. We observed a small, but significant reduction of overall *Xist* RNA levels at day 1 and 2 and a shift towards monoallelic *Xist* expression for mature *Xist* RNA in the mutant compared to the parental cell line (Fig. 6e-f). Pyrosequencing revealed strong skewing towards the inverted allele, which produced 67-84% of *Xist* RNA, pointing to impaired *Xist* upregulation from the chromosome carrying the XertE deletion (Fig. 6g). Through a series of perturbations of the XertE region we could thus confirm that it functions as an enhancer cluster controlling *Xist* transcription in *cis* in an orientation-independent manner. Whether its activity is indeed modulated by *Xert* transcription or whether XertE and the *Xert* transcript function through independent mechanisms remains to be addressed.

### Positive feedback and feedforward loops might amplify *Xert* enhancer activity

Since XertE lies within the *Xert* gene, we asked whether it might affect *Xert* transcription in addition to regulating *Xist*. We thus analyzed *Xert* expression upon XertE perturbation (Fig. 7a-e, Supplementary Fig. 7a-b). We observed a reduction of *Xert* levels upon repression of XertE by CRISPRi (Fig. 7a) and reduced expression from the deleted allele in the ΔXertE mutant mESCs, which was accompanied by a switch from biallelic to monoallelic expression (Fig. 7c-d). Conversely, *Xert* expression was increased when activating XertE with CRISPRa (Supplementary Fig. 7a-b). These findings show that XertE also functions as an enhancer of *Xert* transcription itself. If *Xert* would increase activity of XertE by transcribing through the enhancer cluster, as shown for other lncRNAs (Anderson et al., 2016), such mutual activation could constitute a positive feedback loop between XertE and XertP to amplify *Xist* activation.

**Figure 7.**
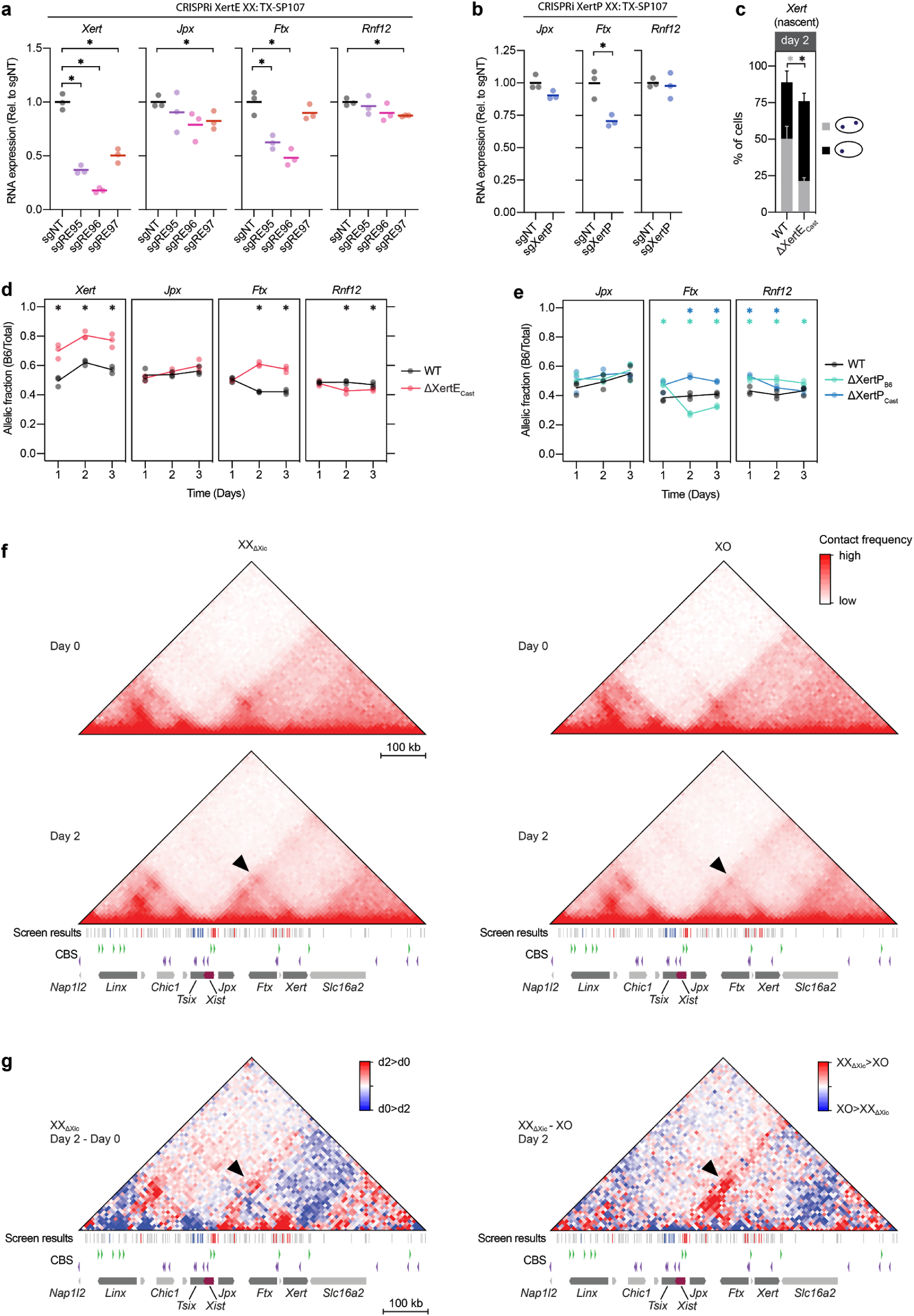
*Xert* and *Ftx* form a regulatory hub that has increased contacts with the *Xist* promoter during initiation of XCI. (**a-b**) CRISPRi repression of individual REs at XertE (a) and of XertP (b) as in Fig. 5b-c and 6b-c. Relative expression of *Xert, Jpx, Ftx and Rnf12* was measured by RT-qPCR (n=3) and normalized to sgNT. (**c-e**) Analysis of Xist regulators in ΔXertE (c-d) and ΔXertP (e) mutant lines by RNA FISH (c) and pyrosequencing (d-e) compared to the parental line (WT). (**f**) Capture-HiC in XX_ΔXic_ (left) and XO cells (right) in the naive state (Day 0, top) and during differentiation (Day 2, bottom). (**g**) Subtraction heatmap of the data shown in (f) for comparing day 0 and 2 in XX_ΔXic_ cells (left) and XX_ΔXic_ and XO cells at day 2 (right). Replicates were merged in (f-g). CTCF binding sites (CBS) and their orientation are indicated below as triangles. The screen results are shown below the tracks, where candidate REs that inhibit (blue) or activate (red) *Xist* expression in the negative or high fractions of the CRISPR screen are colored. Arrowheads in (f) and (g) indicate contacts between Xert and Xist. In (a,b,d,e) horizontal bars denote the mean of 3 biological replicates, dots represent individual measurements. In (c) mean and s.d. of 3 biological replicates are shown. Asterisks indicate significance of *p*<0.05 using an unpaired two-tailed T-test. Colored asterisks in (e) denote comparison of the respective mutant line with the wildtype control.

Next we asked if and how *Xert* might cooperate with other Xist-activators in TAD-E to jointly promote *Xist* upregulation. For this, we analyzed how perturbation of XertE or XertP affected *Jpx*, *Ftx* and *Rnf12*. We found that *Ftx* expression was reduced when targeting XertE or XertP with CRISPRi and increased upon ectopic activation by CRISPRa (Fig. 7a-b, Supplementary Fig. 7a-c). Similarly, *Ftx* showed a clear skewing towards the non-deleted allele in ΔXertP and ΔXertE cells (Fig. 7d-e). These findings show that, in addition to activating *Xist*, the *Xert* elements also promote *Ftx* expression in *cis*. As all *Xert* elements enhance expression of *Ftx*, a well characterized *Xist* activator, *Xert* could in principle enhance *Xist* transcription in an indirect manner via *Ftx*. However, the fact that RE96 in XertE exhibited a stronger effect in the screen than any element within *Ftx* suggests that *Xert*, at least in part, enhances *Xist* transcription independently of *Ftx*.

### Physical contacts between *Xert* and *Xist* are strengthened during *Xist* upregulation

To further corroborate a direct role of *Xert* in *Xist* regulation, we investigated whether the distal *Xert* region would spatially interact with the *Xist* promoter and how such interactions might change during *Xist* upregulation. We performed capture Hi-C (cHi-C) for the *Xic* within our XX_ΔXic_-XO cell model at day 0 and day 2 of differentiation (Fig. 7f). In naive cells, we observed the characteristic split of the *Xic* into TAD-D and TAD-E (Nora et al., 2012) in both cell lines (Fig. 7f, top). During differentiation, a sub-TAD formed within TAD-E, which stretched from the *Xist* promoter to a CTCF-site within *Xert*, thus covering the entire ∼200 kb activating region upstream of *Xist* (Fig. 7f, Supplementary Fig. 7d). A quantitative comparison between naive and differentiating cells revealed an increase in the contact frequency between *Xert* and *Xist* upon differentiation (Fig. 7g, left, Supplementary Fig. 7e, left).

To investigate whether the identified contact patterns were specific for the inactive X, we compared the contact maps between XX_ΔXic_ and XO cells at day 2 of differentiation (Fig. 7g, Supplementary Fig. 7e, right). Contact frequencies of *Xist* with *Xert* and *Ftx* were increased in XX_ΔXic_ compared to XO cells, which might be either a cause or a consequence of *Xist* expression (Fig. 7g, right). In summary, we show that the *Xert* region interacts with *Xist*, supporting its role as an *Xist* enhancer. Moreover, their contact frequency is modulated by differentiation cues and X-chromosomal dosage. Changes in chromatin conformation of the locus might thus contribute to female-specific and mono-allelic *Xist* upregulation at the onset of differentiation.

## Discussion

In the present study, we show how an important developmental locus decodes complex input signals to precisely control gene expression. We identify REs that regulate *Xist* during the onset of random XCI. We then categorize them through chromatin profiling in a cell model that allows dissection of X-dosage sensing and developmental regulation. Hereby we show that only the *Xist* promoter-proximal region responds to X-dosage, while developmental cues activate a ∼200 kb region upstream of *Xist*, containing *Jpx*, *Ftx*, and the newly identified *Xert* region. Through a series of (epi)genome editing approaches we show that the *Xert* promoter (XertP) and a cluster of intronic enhancers (XertE) within *Xert’s* gene body activate *Xist* expression in *cis* and form a regulatory hub with *Ftx*. We can now draw a detailed picture of how distinct transcription factors controlled by X-dosage and differentiation activate specific regulatory regions within the *Xic* to ensure *Xist* upregulation at the primed pluripotent state in a female-specific manner.

We discovered a strong distal enhancer cluster of *Xist*, associated with a previously unknown transcript, which we named *Xert*. It had long been suspected that long-range REs must exist in that region, since a ∼450 kb single-copy transgene containing *Xist* and ∼100 kb of upstream sequence, which includes *Jpx*, but not *Xert* and the *Ftx* promoter, cannot drive *Xist* upregulation in tissues undergoing random XCI *in vivo* or *in vitro (Heard et al., 1996, 1999)*. Since deletion of *Ftx* alone is still compatible with female development (Soma et al., 2014), we suggest that *Ftx* and *Xert* together form a regulatory hub, wherein their transcripts and enhancer elements promote each other’s activity to jointly allow strong *Xist* upregulation upon differentiation. For both, *Xert* and *Ftx*, their strongest REs (RE85/96) lie within their major transcripts. At both loci transcription might help to activate transcript-embedded enhancers, as shown previously at the *Hand2* locus (Anderson et al., 2016). Since nascent transcription can block H3K27me3 deposition (Hosogane et al., 2016; Kaneko et al., 2014; Laugesen et al., 2019), transcription might also accelerate removal of the repressive H3K27me3 hotspot, which covers the entire region before differentiation and is cleared more rapidly from the transcribed loci compared to their surroundings (see Fig. 3b). While *Ftx* is expressed rather ubiquitously (Chureau et al., 2011), *Xert* transcription appears to be restricted to a short period when random XCI is initiated. This transient activation of the Xert region might explain why a GFP reporter inserted downstream of the *Xist* promoter was found to be only transiently expressed at the onset of differentiation (Loos et al., 2016). Since *Xert* seems to primarily boost *Xist* expression levels, as revealed by the binned sorting strategy we used in our CRISPR screen, its activation at the onset of XCI might be important to pass a previously postulated activation threshold (Monkhorst et al., 2008; Mutzel and Schulz, 2020; Mutzel et al., 2019). Subsequent *Xert* downregulation might help to prevent spurious *Xist* upregulation from the Xa, while *Ftx* and *Jpx* maintain *Xist* expression on the Xi in somatic cells.

Our results finally answer the long-standing question of how developmental regulation of *Xist* is ensured. We show that, in addition to downregulation of the repressors *Tsix* and REX1, *Xist* upregulation requires activation of a series of distal enhancer elements, which appear to be controlled by primed pluripotency factors. Among these are SMAD2/3, which are activated by the TGFβ/activin pathway. The activin receptor has previously been identified as XCI activator in two different shRNA screens, further supporting a role of this pathway in *Xist* regulation (Bhatnagar et al., 2014; Sripathy et al., 2017). Intriguingly, the TGFβ pathway is also regulated by RNF12, which enhances SMAD2/3 signaling via degradation of inhibitory SMAD7 (Zhang et al., 2012). This might be the reason why *Xert* is transcribed slightly more than double in cells with two X chromosomes, which might also contribute to X-dosage dependent *Xist* regulation. Nevertheless, the distal enhancer elements in the *Ftx-Xert* region were strongly activated both in XX and XO cells upon differentiation, showing that they mainly sense developmental progression.

X-dosage, by contrast, primarily acts on *Xist’s* promoter-proximal region, including a CpG island ∼1.5 kb downstream of the TSS and a region encoding the repeat A of the *Xist* RNA, both of which have previously been implicated in *Xist* regulation (Hoki et al., 2009; McDonald et al., 1998; Norris et al., 1994; Royce-Tolland et al., 2010) (see Supplementary screen discussion). The region is bound by CTCF, YY1 and REX1 (Makhlouf et al., 2014; Navarro et al., 2006), with REX1 being targeted for degradation in an X-dosage dependent manner (Gontan et al., 2018), further supporting a role of this region in X-dosage sensing. YY1 and also CTCF, both of which have been implicated in long-range chromatin interactions (Nora et al., 2017; Weintraub et al., 2017), bind this region preferably on the Xi in somatic cells, with binding to the Xa likely being inhibited by DNA methylation (Calabrese et al., 2012; Chapman et al., 2014; Makhlouf et al., 2014; Norris et al., 1994). Differential CTCF and YY1 binding between the alleles might underlie the increase in long-range contacts that we observe on the *Xist*-expressing chromosome. At the same time when the *Xist* promoter is activated on the future Xi, we observe active repression at the Xa through deposition of H3K9me3. This might be mediated by TRIM28/KAP1, which has been reported to bind the region on the Xa (Enervald et al., 2020) and recruits H3K9-specific histone methyl-transferases (Ecco et al., 2017). How KAP1 is targeted to the region however remains an open question.

Overall our analyses reveal that the *Xic* assumes at least three distinct states (Fig. 8). In undifferentiated mESCs, the *Xist* promoter is accessible, but transcription is repressed by *Tsix* and REX1, while distal enhancers are repressed by the H3K27me3 hotspot. Upon differentiation, distal enhancers are derepressed and activated by primed pluripotency factors, resulting in upregulation of *Jpx*, *Ftx* and *Xert*. Those distal regions will then drive *Xist* upregulation, but only if the promoter-proximal region is maintained in an active configuration by X-dosage dependent mechanisms, thereby restricting *Xist* upregulation to females. In males, and presumably also on the future Xa in females, the promoter region assumes a heterochromatic state. Activation by distal enhancers and active repression thus appear to be two competing processes at the *Xist* promoter and their relative dynamics must be tightly tuned in an X-dosage dependent manner.

**Figure 8.**
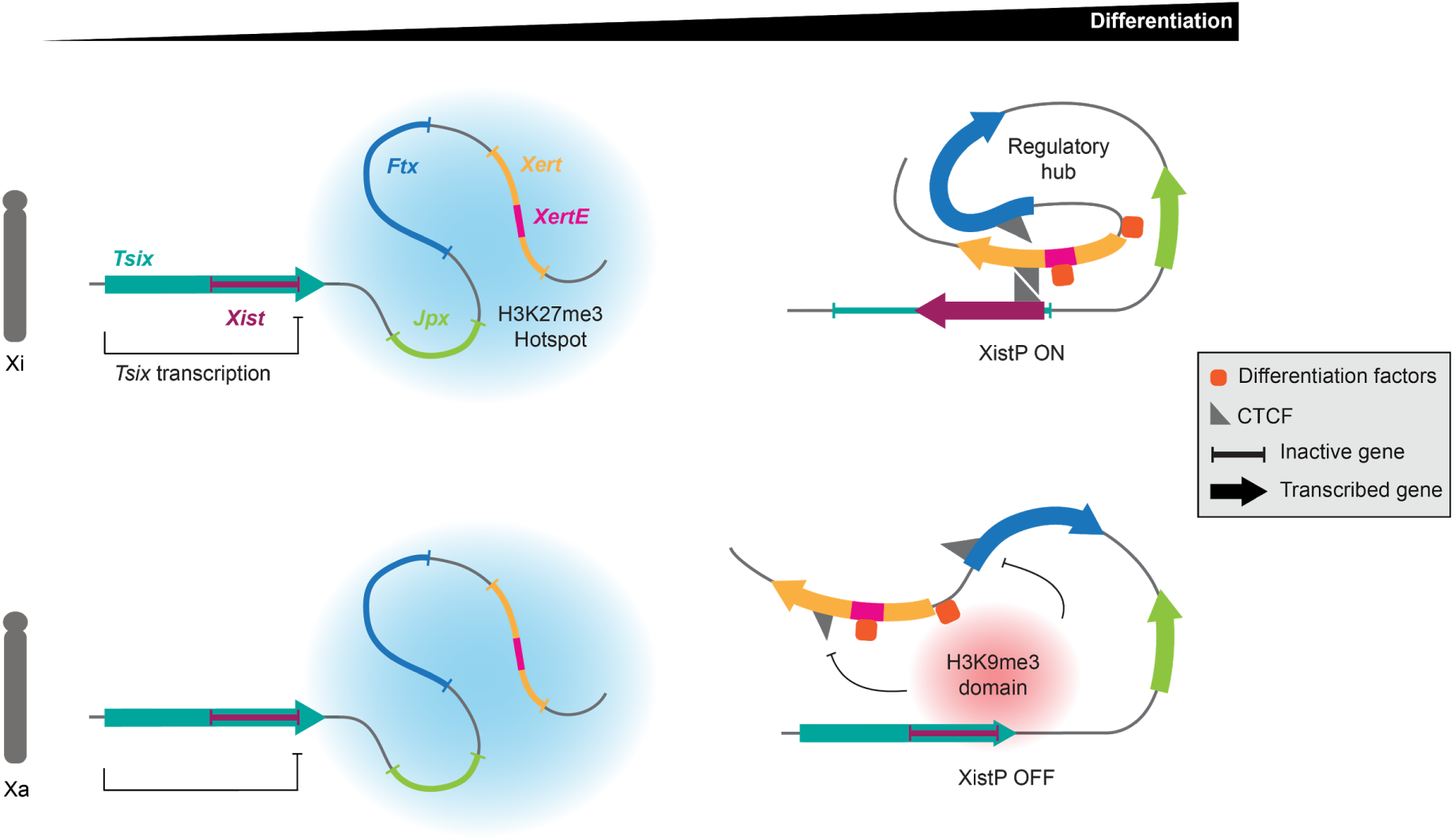
A model of initial *Xist* upregulation. Schematic of *Xist* regulation where *Xist* activators *Jpx*, *Ftx* and *Xert* are repressed in undifferentiated cells, in part by a H3K27me3 hotspot (blue cloud), and *Xist* is repressed by antisense transcription through *Tsix*, on the Xi and on the Xa. Following differentiation, *Xist* activators are upregulated and their associated enhancers are activated by differentiation factors. The *Xist* promoter (XistP) acquires a repressive H3K9me3 domain (red cloud) at the Xa, which prevents *Xist* contacts with *Xert* and *Ftx*. At the Xi, XistP is in an active configuration allowing CTCF recruitment and contacts with distal activating elements thereby inducing *Xist* upregulation.

Taken together, we have uncovered a regulatory hierachy at the *Xic*, which allows coincidence detection of two signals that inform the locus on sex and developmental stage of the cell. Similar to other developmental genes, multiple distal elements function as tissue-specific enhancers. The promoter-proximal region by contrast acts as a binary switch, which, when turned off, renders the core promoter unresponsive to long-range regulation. In this way, two signals controlling distal and proximal elements, respectively, are integrated with an AND-logic. Our findings are thus the first step towards understanding how logical operations are performed by *cis*-regulatory landscapes to generate the complex expression patterns of developmental genes in mammals.

## Methods

### Cell lines

The female TX1072 cell line (clone A3) is a F1 hybrid ESC line derived from a cross between the 57BL/6 (B6) and CAST/EiJ (Cast) mouse strains that carries a doxycycline-responsive promoter in front of the *Xist* gene on the B6 chromosome and an rtTA insertion in the *Rosa26* locus (Schulz et al., 2014). TXΔXic_B6_ (here referred to as XX_ΔXic_) carries a 773 kb deletion around the *Xist* locus on the B6 allele (chrX:103,182,701-103,955,531, mm10) (Pacini et al., 2020). The TX1072 XO line (clone B7) has lost the B6 X chromosome and is trisomic for chromosome 16. Female 1.8 XX mESCs carry a homozygous insertion of 7xMS2 repeats in *Xist* exon 7 and are a gift from the Gribnau lab (Schulz et al., 2014). The female TXΔXertP (Clone B5 and D5), TXΔXertE (Clone F6) and TX-Xert-pA (Clone C4 and F11) cell lines were generated by introducing a heterozygous deletion or insertion of a polyA cassette with a puromycin selectable marker (Supplementary Fig. 5i) in TX1072 mESCs. The B6 chromosome is modified in TXΔXertP B5 and both TX-Xert-pA lines, and the Cast allele carries the deletion in TXΔXertP D5 and TXΔXertE. The cell lines were generated using CRISPR-Cas9 mediated genome editing (see below) and the deleted regions are specified in Supplementary Table 6. The TXΔXertE line also carries an inversion on the B6 allele, TXΔXertP B5 carries duplications of parts of Chr 10 and TXΔXertP D5 is trisomic for Chr 8 (Supplementary Fig. 5f).

The male E14-STN_ΔTsixP_ mESC cell line expresses the CRISPRa Sun-Tag system (Tanenbaum et al., 2014) under a doxycycline-inducible promoter and carries a 4.2 kb deletion around the major *Tsix* promoter (ChrX:103445995-103450163, mm10, Supplementary Table 6). The cell line was generated by introducing the *Tsix* deletion in E14-STN mESCs (Heurtier et al., 2019) (a kind gift from Navarro lab) and NGS karyotyping (see below) detected duplications of parts of Chr 2.

The TX-SP106 (Clone D5) mESC line stably expresses PYL1-VPR-IRES-Blast and ABI-tagBFP-SpdCas9, constituting a two-component CRISPRa system, where dCas9 and the VPR activating domain are fused to ABI and PYL1 proteins, respectively, which dimerize upon treatment with abscisic acid (ABA). The TX-SP107 (Clone B6) mESC line stably expresses PYL1-KRAB-IRES-Blast and ABI-tagBFP-SpdCas9, constituting a two-component CRISPRi system, where dCas9 and the KRAB repressor domain are fused to ABI and PYL1 proteins, respectively, which dimerize upon ABA treatment. Both cell lines were generated through piggybac transposition (see below). Correct karyotype was confirmed for TX-SP106 (Clone D5) and TX-SP107 (Clone B6) by NGS (Supplementary Fig. 5f). T Since repression in TX-SP107 cells transduced with sgRNAs was often observed already without ABA treatment, we could not make use of the inducibility of the system. Instead, TX-SP107 cells were always treated with ABA (100 µM) 24 h before the analysis and effects were compared to NTC sgRNAs.

#### mESC culture and differentiation

TX1072 mESCs, TX1072 derived mutant cell lines and 1.8 cells were grown on 0.1% gelatin-coated flasks in serum-containing medium supplemented with 2i and LIF (2iL) (DMEM (Sigma), 15% ESC-grade FBS (Gibco), 0.1 mM β-mercaptoethanol, 1000 U/ml leukemia inhibitory factor (LIF, Millipore), 3 μM Gsk3 inhibitor CT-99021, 1 μM MEK inhibitor PD0325901, Axon). Differentiation was induced by 2iL withdrawal in DMEM supplemented with 10% FBS and 0.1 mM β-mercaptoethanol at a density of 1.6*10^4^ cells/cm^2^ on fibronectin-coated (10 μg/ml) tissue culture plates, if not stated otherwise. During the pooled CRISPR screen and CRISPRi experiments, cells were differentiated at a density of 3.6*10^4^ cells/cm^2^. For STARR-seq, 1*10^5^ cells/cm^2^ cells were seeded for 2iL conditions, while 7*10^4^ cells/cm^2^ were used for differentiation. E14-STN_ΔTsixP_ mESC cells were grown on 0.1% gelatin-coated flasks in serum-containing medium (DMEM (Sigma), 15% ESC-grade FBS (Gibco), 0.1 mM β-mercaptoethanol), supplemented with 1000 U/ml LIF (SL). Differentiation was induced by LIF withdrawal in DMEM supplemented with 10% FBS and 0.1 mM β-mercaptoethanol at a density of 5.2*10^4^ cells/cm^2^ in fibronectin-coated (10 μg/ml) tissue culture plates.

### Molecular Cloning

#### Cloning sgRNA plasmids

For genomic deletions, sgRNAs were designed to target the 5’ and 3’ end of the region of interest and cloned into pSpCas9(BB)-2A-Puro (pX459) V2.0 (Ran et al., 2013). For the pA-insertion lines a sgRNA that targeted a site downstream of the XertP was cloned into pX330-U6-Chimeric_BB-CBh-hSpCas9 (pX330) (Cong et al., 2013). pX459 and pX330 were kind gifts from Feng Zhang (Addgene plasmid # 42230 and # 62988). SgRNAs (sequences are given in Supplementary Table 6) were cloned following the Zhang lab protocol (https://media.addgene.org/cms/filer_public/6d/d8/6dd83407-3b07-47db-8adb-4fada30bde8a/zhang-lab-general-cloning-protocol-target-sequencing_1.pdf). In short, two complementary oligos containing the guide sequence and a BbsI recognition site were annealed and ligated with the BbsI (New England Biolabs) digested target plasmid. The ligation mixes were heat shock transformed into NEB Stable competent cells (New England Biolabs) and grown as single colonies on LB-Agar plates (supplemented with Ampicillin 100 ug/ml) overnight at 37 °C. Single colonies were expanded and confirmed with Sanger sequencing.

#### pA insertion cassette plasmid

Left and right homology arms of ∼500 bp, flanking the Cas9 cut site, were amplified from TX1072 genomic DNA with Q5®High-Fidelity DNA Polymerase (New England Biolabs) and restriction sites for NdeI/XhoI and EcoRI/PmeI were included in the 5’ and 3’ primers (PK47, PK64, PK65, PK66, Supplementary Table 6). To prevent Cas9 cutting the homology arm following integration of the pA cassette, a point mutation of the PAM was introduced in the cloning primers. The homology arms were digested with their corresponding restriction enzymes, gel-purified and cloned into pFX5 (Galupa et al., 2020) (a kind gift from Edith Heard) that contains a puromycin selectable pA cassette between the site-specific homology arms, resulting in plasmid SP292. For each round of cloning, the ligation mixes were heat-shock transformed into NEB Stable competent cells (New England Biolabs) and grown as single colonies on LB-Agar (supplemented with Ampicillin 100 μg/ml) plates overnight at 37 °C. Single colonies were expanded and confirmed with Sanger sequencing.

#### Cloning of sgRNAs in multiguide expression system

For CRISPRa and CRISPRi three different sgRNAs targeting the same RE (Supplementary Table 6) were cloned into a single sgRNA expression plasmid with Golden Gate cloning, such that each sgRNA was controlled by a different Pol III promoter (mU6, hH1 or hU6) and fused to the optimized sgRNA constant region described in Chen et al (Chen et al., 2013). To this end, the sgRNA constant region of the lentiGuide-puro sgRNA expression plasmid (Sanjana et al., 2014) (Addgene 52963) was exchanged for the optimized sgRNA constant region, thus generating the vector SP199. The vector was digested with BsmBI (New England Biolabs) overnight at 37 °C and gel-purified. Two fragments were synthesized as gene blocks (IDT) containing the optimized sgRNA constant region coupled to the mU6 or hH1 promoter sequences. These fragments were then amplified with primers that contained part of the sgRNA sequence and a BsmBI restriction site (primer sequences can be found in Supplementary Table 6) and purified using the gel and PCR purification kit (Macherey & Nagel). The vector (100 ng) and two fragments were ligated in an equimolar ratio in a Golden Gate reaction with T4 ligase and the BsmBI isoschizomer Esp3I for 20 cycles (5 min 37 °C, 5 min 20 °C) with a final denaturation step at 65 °C for 20 min. Vectors were transformed into NEB Stable competent E.coli. Successful assembly was verified by ApaI digest and Sanger sequencing.

### Piggybac transposition

TX-SP106 and TX-SP107 lines were generated by piggybac transposition. To this end the puromycin resistance cassette in the piggybac CRISPRa and CRISPRi expression plasmid (pSLQ2817 and pSLQ2818) was exchanged for a blasticidin resistance, resulting in plasmid SP106 and SP107 respectively. pSLQ2817 and pSLQ2818 were gifts from Stanley Qi (Gao et al., 2016) (Addgene plasmids #84239 and #84241). The respective plasmid was then transfected with Lipofectamin 2000 (Thermo Fisher Scientific) into female TX1072 mESCs in a 5-to-1 transposase-to-target ratio with the hyperactive transposase (pBroad3_hyPBase_IRES_tagRFP) (Redolfi et al., 2019). RFP-positive cells were sorted 24 h after transfection and expanded as single clones under blasticidin selection (5 ng/µl, Roth).

### Genome Engineering

#### Generation of promoter/enhancer deletion and pA insertion mESC lines

To generate deletions up to 4*10^6^ TX1072 (ΔXertP, ΔXertE) or E14-STN (ΔTsixP) mESCs, cultured in gelatin-coated flasks in SL medium, were nucleofected with 2 µg of each sgRNA/Cas9 plasmid and 3 pmol or 30 pmol (for E14-STN_ΔTsixP_) repair oligo (Supplementary Table 6) using the Lonza 4D-Nucleofector with the P3 Primary Cell 4D-NucleofectorTM kit (Lonza) and CP-106 (for TXΔXertP and E14-STN_ΔTsixP_) or CG-104 (for TXΔXertE) nucleofection programs. Afterwards the cells were plated on gelatin-coated 10 cm plates with SL medium. Between 18 and 24 h following nucleofection, cells were selected in SL medium supplemented with puromycin (1 ng/ml) for 24 h. Two to 3 days later, the cells were trypsinized and seeded at low densities in gelatin-coated 10 cm plates in SL medium wherein they were cultured until single colonies were visible (up to 12 days).

In order to insert the pA cassette downstream of the *Xert* TSS, 4*10^5^ TX1072 cells cultured in SL medium were reverse transfected with 250 ng pX330-PK45/46 and 750 ng of the pA cassette-containing donor plasmid (SP292) with Lipofectamine 3000 reagent using 1 µl P3000 reagent and 6 µl Lipofectamine 3000 (Thermo Fisher Scientific) following the manufacturer’s guidelines. Cells were incubated with the lipofection mix for 10 min at room temperature (RT) and then seeded into a gelatin-coated 12-well plate in 1 ml of SL medium. Medium was exchanged daily. Two days after transfection, cells were split to low density in gelatin-coated 10 cm plates. The following day, medium was exchanged for SL medium with puromycin (1 ng/ml) to select for cells carrying the insertion, selection medium was exchanged every second day until colonies appeared. Single colonies were manually picked, trypsinized in a 96-wells plate, subsequently transferred to gelatin-coated 96-wells plates and cultured to semi-confluency in SL.

#### Genotyping of engineered clones

Semi-confluent 96-well plates with clones were split into 2 low density and 1 high density gelatin-coated 96-well plates with SL medium. Up to 2 days later gDNA was isolated from the high density plate. The cells were washed with PBS and lysed in the 96-wells plate with 50 µl Bradley lysis buffer (10 mM Tris-HCl pH 7.5, 10 mM EDTA, 0.5% SDS, 10 mM NaCl, 1 mg/ml Proteinase K (Invitrogen)). The plate was incubated overnight at 55 °C in a humidified chamber. To precipitate gDNA, 150 µl ice-cold 75 mM NaCl in 99% EtOH was added per well and the plate was incubated for 30 min at RT. The plate was centrifuged for 15 min at 4000 rpm and 4 °C. The pellet was washed once with 70% EtOH and centrifuged for 15 min at 4000 rpm and air dried at 45 °C for 10 min. The gDNA was resuspended in 150 µl TE buffer (10 mM Tris pH 8.0 and 1 mM EDTA, pH 7.5) for 1 h at 37 °C. The clones were initially characterized by PCR using either Qiagen HotStarTaq Plus kit (Qiagen) or Q5 High Fidelity DNA polymerase (New England Biolabs) following the manufacturer’s guidelines, and primer combinations that distinguish between WT and deletions, insertions, or inversions (Supplementary Table 6). A small number of positive clones were expanded from low density plates. PCR genotyping was repeated on gDNA isolated using the DNeasy Blood and Tissue Kit (Qiagen). To identify the targeted allele, amplicons containing SNPs were gel-purified and sequenced. Primers and SNP position are given in Supplementary Table 6. Few clones were selected and adapted to 2iL medium for at least 4 passages prior to subsequent experiments. For E14-STN_ΔTsixP_, clone C6 was further sub-cloned and following PCR genotyping (Supplementary Fig. 5a-b), subclone E14-STN_ΔTsixP_ B2 was chosen for future experiments (here referred to as E14-STN_ΔTsixP_).

### NGS Karyotyping

Cell lines were karyotyped via double digest genotyping-by-sequencing (ddGBS), a reduced representation genotyping method, as described previously (Genolet et al., 2020). Briefly, the forward and reverse strands of a barcode adapter and common adapter were diluted and annealed, after which they were pipetted into each well of a 96-well PCR plate together with 1 μg of each sample and dried overnight (oligo sequences are listed in Supplementary Table 6). The following day samples were digested with 20 μl of a NIaIII and PstI enzyme mix (New England Biolabs) in NEB Cutsmart Buffer at 37 °C for 2 h. After the digest, a 30 μl mix with 1.6 μl of T4 DNA ligase (New England Biolabs) was added to each well and placed on a thermocycler (16 °C 60 min followed by 80 °C 30 min for enzyme inactivation). By doing this, barcode and common adapters with ends complementary to those generated by the two restriction enzymes were ligated to the genomic DNA.

Samples were cleaned with CleanNGS beads (CleanNA) using 90 μl of beads for each well and following manufacturers instructions. Samples were eluted in 25 μl ddH_2_O and DNA was quantified using a dsDNA HS Qubit assay (Thermofisher). Samples were pooled in an equimolar fashion, size-selected (300-450bp) by loading 400 ng of each pooled sample on an agarose gel followed by a cleaning step using the Nucleospin Gel and PCR Cleanup kit (Macherey-Nagel). Samples were PCR amplified using the Phusion High-Fidelity DNA Polymerase (New England Biolabs) and an annealing temperature of 68 °C over 15 amplification cycles (OG218/OG219). Resulting amplicons were cleaned with CleanNGS beads in a 1:1.2 ratio (sample:beads) and sequenced with 2×75bp on the Miseq platform (12 pM loading concentration), yielding up to 0.2*10^6^ fragments per sample.

Data processing and statistical analysis was performed on the public Galaxy server usegalaxy.eu. Fastq files were uploaded and demultiplex using the “Je-demultiplex” tool (Girardot et al. 2016). Reads of the karyotyping analysis were mapped to the mouse genome (mm10) using BWA (Li and Durbin, 2009). The reads for each chromosome were then counted using deeptols(Ramírez et al., 2016) v2.0) on the useGalaxy platform(Afgan et al., 2016; Giardine et al., 2005) with option [multiBamSummary]. The counts per chromosome were divided by the sum of all counts per sample. The relative counts were then normalized to the wildtype and visualized as a heatmap (Supplementary Fig. 5f).

### RNA extraction, reverse transcription, qPCR

Cells were lysed directly in the plate by adding up to 1 ml of Trizol (Invitrogen). RNA was isolated using the Direct-Zol RNA Miniprep Kit (Zymo Research) following the manufacturer’s instructions with on-column DNAse digestion. If cDNA was subsequently analyzed by pyrosequencing, DNase digest was performed using Turbo DNA free kit (Ambion). Up to 1 µg RNA was reverse transcribed using Superscript III Reverse Transcriptase (Invitrogen) with random hexamer primers and expression levels were quantified in the QuantStudio™ 7 Flex Real-Time PCR machine (Thermo Fisher Scientific) using Power SYBR™ Green PCR Master Mix (Thermo Fisher Scientific) normalizing to Rrm2 and Arpo. Primers used are listed in Supplementary Table 6.

### 3’- and 5’RACE

To identify transcript isoforms as well as exact stop and start sites of *Xert*, 3’- and 5’RACE were performed. First, RNA was isolated from 2 day-differentiated TXΔXic_B6_ cells using the Direct-Zol RNA Miniprep Kit (Zymo Research). To remove any remaining gDNA, RNA samples were rigorously treated with DNase for 20 min at 37 °C using the TURBO DNA-freeTM Kit (Thermo Fisher Scientific) according to the manufacturer’s instructions. Poly-adenylated RNAs were purified from 5 µg total RNA with the Dynabeads®Oligo (dT)25 Kit (Thermo Fisher Scientific) following the manufacturer’s instructions.

For 3’-RACE cDNA was synthesised as described before, instead using 50 ng purified polyadenylated RNA and the oligo(dT)-anchor primer from the 5’/3’RACE kit, by following the manufacturer’s guidelines. To remove DNA:RNA duplexes, 25 ng of cDNA was digested with 0.5 µl 1:40 diluted RNaseH (New England Biolabs) for 20 min at 37 °C. To specifically amplify the 3’ end of the transcript for 3’RACE, RNaseH-treated cDNA was PCR-amplified by Phusion High-Fidelity DNA Polymerase (Thermo Fisher Scientific) according to the manufacturer’s instructions using the gene-specific forward primer PK1 and the anchor primer PK9. PCR products were analysed on agarose gels and purified using QIAquick PCR Purification Kit (Qiagen) according to the manufacturer’s instructions. To increase specificity the isolated PCR product was PCR amplified with the nested gene-specific forward primer PK4 and the anchor primer PK9. Additionally, a PCR targeting putative exon 2 and exon 6, was performed using the gene-specific PK4 and PK17 primers.

For 5’-RACE, 50 ng of purified poly-adenylated RNA was reverse transcribed using the gene-specific reverse primer PK35. To remove DNA:RNA duplexes, cDNA was digested with RNaseH as described before. RNaseH-treated cDNA was purified using QIAquick PCR purification Kit according to the manufacturer’s instructions. Subsequently, 5’ pA tailing of the product was performed with the 5’/3’RACE kit, 2nd generation (Roche) according to the manufacturer’s guidelines. The anchor sequence was added to the 5’ end of the transcript by PCR amplification using the gene-specific reverse primer PK13 and the oligo(dT)-anchor primer and Phusion High-Fidelity DNA Polymerase (New England Biolabs). PCR products were analysed on agarose gels and purified using the QIAquick PCR purification Kit according to the manufacturer’s instructions. To increase specificity, the cleaned PCR product was amplified in a nested PCR using the nested gene-specific reverse primer PK34 and the anchor primer PK9. All primer sequences are given in Supplementary Table 6.

#### TOPO TA Cloning and Sanger Sequencing

Blunt-end PCR amplicons underwent A-tailing using HotStarTaq DNA Polymerase (New England Biolabs). The PCR products from the nested 3’/5’RACE were cleaned-up using QIAquick PCR purification Kit and then mixed with 5 µl 10x DNA polymerase reaction buffer, 10 µl of 1 mM dATP, 0.2 µl of HotStarTaq DNA polymerase, filled up to 50 µl with nuclease free water and incubated for 20 min at 72 °C. The A-tailed PCR products were separated on agarose gels, and bands were individually isolated from agarose gels (Supplementary Fig. 4c) using the QIAquick Gel Extraction Kit (Qiagen) and cloned into TOPO vector pCR2.1 using the Topo TA cloning kit (Invitrogen) according to the manufacturer’s instructions. For the ligation, 1 µl 10x T4 DNA ligase buffer, 1.5 µl pCR2.1 vector, 10 ng A-tailed gel-isolated PCR product and 1 µl T4 DNA ligase (New England Biolabs) were mixed in a total reaction volume of 10 µl and incubated for 15 min at room temperature. One Shot® TOP10 chemically competent E. coli (Thermo Fisher Scientific) were heat shock transformed and plated on LB-agar plates supplemented with Ampicillin 100ug/ml, 100 µl 20 mg/ml X-gal (Sigma Aldrich). Plates were incubated overnight at 37 °C, the following day colonies were assessed by blue/white screening. Five white colonies were picked per plate, inoculated in 5 ml LB medium (supplemented with Ampicillin 100 μg/ml) and shaken overnight at 37 °C. Plasmids were purified from the bacterial cultures using Plasmid Mini Prep (PeqLab) according to the manufacturer’s instructions and analyzed by Sanger sequencing via LGC Genomics GmbH, PK11 was used as sequencing primer. The obtained sequence data between the gene-specific forward primer (for 3’RACE) or gene-specific reverse primer (for 5’-RACE) and the anchor primer was extracted and aligned to the mouse genome (mm10) via basic local alignment search tool (BLAST) and visualized using the UCSC genome browser (Supplementary Fig. 4d).

### Poly-adenylated RNA-seq and *de novo* transcriptome assembly

Total RNA (100ng) from 2 days differentiated TX1072 mESCs was subjected to strand-specific RNA-seq library preparation with the TruSeq® RNA Sample Preparation Kit v2 (Illumina), which included polyadenylated RNA enrichment using oligo-dT magnetic beads, by following the manufacturers guidelines. The libraries were subjected to Illumina NGS PE50 on the HiSeq 4000 platform to obtain approximately 65 million fragments. Reads were aligned using *STAR* (v2.7.5a) (Dobin et al., 2013) with options [--outSAMattributes NH HI NM MD]. For de novo transcript assembly, the sorted bam file was analyzed in *Cufflinks* (v2.2.1) with the parameter [--library-type fr-firststrand]. Mapping statistics can be found in Supplementary Table 4.

### Xert isoforms detection and analysis

3’ and 5’RACE had identified multiple isoforms of *Xert* with a length of 398-767bp (Supplementary Table 5). To verify the *Xert* isoforms as detected by 3’/5’-RACE, we generated a Sashimi plot in IGV (v2.3.94) and analyzed the transcripts predicted by the *de novo* transcript assembly based on polyadenylated RNA-seq data (see above) (Supplementary Fig. 4c). These two analyses indicated two additional unidentified *Xert* isoforms, which were confirmed by conventional PCR with primers PK4+PK17 on RNaseH treated cDNA as described above. Sanger sequencing of isolated bands A1-4 (Supplementary Fig. 4b) indeed revealed these 2 additional *Xert* isoforms (Supplementary Fig. 4c).

Our 5’-RACE data suggested that *Xert* contains 2 TSSs. However, since we detected band D6 (Supplementary Fig. 4c) only once from all 5’-RACE colonies analyzed, and this far upstream TSS was neither detected in RNA-seq nor in TT-seq data (Fig 4a), we considered the TSS starting at ChrX:103637012 (5’-RACE bands D1-5 in Supplementary Fig. 4c) the only TSS driving *Xert* transcription in our mESCs.

To detect any open reading frames (ORFs) that could potentially code for protein, DNA sequences from all processed *Xert* transcript isoforms were loaded into Geneious (v10.2.6) and assessed with the Find ORF option for a minimum size of 150 bp in 5’-3’ direction. Six ORFs with a length between 153bp and 234bp were identified (Supplementary Table 5).

### Pyrosequencing

To quantify relative allelic expression for individual genes, an amplicon containing a SNP at the Cast allele was amplified by PCR from cDNA using Hot Start Taq (Qiagen) for 38 cycles. The PCR product was sequenced using the Pyromark Q24 system (Qiagen). Assay details are given in Supplementary Table 6.

### RNA FISH

RNA FISH was performed using Stellaris FISH probes (Biosearch Technologies). Probe details can be found in Supplementary Table 6. Cells were dissociated using Accutase (Invitrogen) and adsorbed onto coverslips (#1.5, 1 mm) coated with Poly-L-Lysine (Sigma) for 5 min. Cells were fixed with 3% paraformaldehyde in PBS for 10 min at RT (18–24 °C) and permeabilized for 5 min on ice in PBS containing 0.5% Triton X-100 and 2 mM Ribonucleoside Vanadyl complex (New England Biolabs). Coverslips were preserved in 70% EtOH at −20 °C. Prior to FISH, coverslips were incubated for 5 minutes in wash buffer containing 2x SSC and 10% formamide, followed by hybridization for 6 hours to overnight at 37 °C with 250 nM of each FISH probe in 50 μl Stellaris RNA FISH Hybridization Buffer (Biosearch Technologies) containing 10% formamide. Coverslips were washed twice for 30 min at 37 °C with 2x SSC/10% formamide with 0.2 mg/ml Dapi being added to the second wash. Prior to mounting with Vectashield mounting medium coverslips were washed with 2xSSC at RT for 5 minutes. Images were acquired using a widefield Z1 Observer microscope (Zeiss) using a 100x objective.

### Lentiviral transduction

To package lentiviral vectors into lentiviral particles, 1*10^6^ HEK293T cells were seeded into one well of a 6-well plate and transfected the next day with the lentiviral packaging vectors: 1.2 µg pLP1, 0.6 µg pLP2 and 0.4 µg pVSVG (Thermo Fisher Scientific), together with 2 µg of the desired construct using Lipofectamine 2000 (Thermo Fisher Scientific). HEK293T supernatant containing the viral particles was harvested after 48 h. 0.2*10^6^ mESCs were seeded per well in a 12-well plate with 2iL (for TX-SP106 and TX-SP107) or SL medium (for E14-STN_ΔTsixP_) and transduced the next day with 1ml of 5:1 concentrated (lenti-X, Clontech) and filtered viral supernatant with 8 ng/µl polybrene (Sigma Aldrich). Puromycin selection (1 ng/µl, Sigma Aldrich) was started two days after transduction and kept for at least 2 passages.

### Flow-FISH

For Flow-FISH the PrimeFlow RNA assay (Thermofisher) was used according to the manufacturer’s recommendations. Specifically, the assay was performed in conical 96-well plates with 5*10^6^ cells per well with Xist specific probes, labelled with Alexa-Fluor647 (VB1-14258) (Thermo Fisher Scientific). Samples were resuspended in PrimeFlow™ RNA Storage Buffer before flow cytometry. Cells were analyzed or sorted using the BD FACSAria^TM^ II or BD FACSAria™ Fusion flow cytometers. The sideward and forward scatter areas were used for live cell gating, whereas the height and width of the sideward and forward scatter were used for doublet discrimination. At least 20.000 cells were measured per replicate. FCS files were gated using RStudio with the *flowCore* (v1.52.1) and *openCyto* packages (v1.24.0) (Finak et al., 2014; Hahne et al., 2009). All cells that showed a fluorescent intensity above the 99th-percentile of the 2iL-control were marked as *Xist*-positive. These cells were then used to calculate the mean fluorescent intensity in the *Xist*-positive fraction after background correction by subtracting the mean intensity of the 2iL-control. Both, the percentage of *Xist*-positive cells and the mean fluorescent intensity of the *Xist*-positive fraction were plotted as a ratio to the non-targeting control.

### NGS data processing

#### Genome preparation

For all alignment of data generated within the TX1072 cell lines, all SNPs in the mouse genome (mm10) were N-masked (Barros de Andrade E Sousa et al., 2019; Pacini et al., 2020) using *SNPsplit* (v0.3.2) (Krueger and Andrews, 2016) for high-confidence SNPs between present in the TX1072 cell line as described previously (Barros de Andrade E Sousa et al., 2019; Pacini et al., 2020). For all other data (STARR-seq, published data) data was aligned to the reference genome.

#### Read filtering

Following alignment, sequencing data was filtered for mapped and, for paired-end data, properly paired reads using *samtools* (Li et al., 2009) (v1.10) with options [view -f 2 -q 20] for ATAC-seq, CUT&Tag and paired ChIP-seq, [view -F 4 -q 20] for unpaired ChIP-seq, [view -f 2 -q 10] for STARR-seq and [view -q 7 -f 3] for RNA-seq and TT-seq data. Afterwards, the BAM files were sorted using *samtools* (Li et al., 2009) (v1.10) with [sort]. Blacklisted regions for mm10 (ENCODE Project Consortium, 2012) were then removed using *bedtools* (Quinlan and Hall, 2010) (v2.29.2) with options [intersect -v]. Unless stated otherwise, duplicates were marked and removed using *Picard* (v2.18.25) with options [MarkDuplicates VALIDATION_STRINGENCY=LENIENT REMOVE_DUPLICATES=TRUE] (http://broadinstitute.github.io/picard). For analysis, BAM files of individual replicates were merged using samtools (Li et al., 2009) (v1.10) with [merge].

#### Generation of coverage tracks

BIGWIG coverage tracks for sequencing data were created using *deeptools2* (v3.4.1) (Ramírez et al., 2016) merged replicates, if available. For TT-seq and polyadenylated RNA-seq, BAM files were split depending on the strand prior to track generation. Normalization was performed using the total number of autosomal reads. RNA-seq and unpaired ChIP-seq data was processed with the options [bamCoverage -bs 10 --normalizeUsing CPM -ignore chrX chrY]. For paired and unspliced data types (ATAC-seq, CUT&Tag, ChIP-seq & TT-seq) reads were additionally extended using[-e]. The tracks were visualized using the UCSC genome browser (Kent et al., 2002).

#### Peak calling

Unless stated otherwise, peaks were called using *MACS2* (Zhang et al., 2008) (v2.1.2) with standard options [callpeak -f BAMPE/BAM -g mm -q 0.05] on individual replicates. For ChIP-seq, input samples were included for normalization using [-c]. Only peaks detected in all replicates were retained by merging replicates using *bedtools* (Quinlan and Hall, 2010) (v2.29.2) with [intersect].

### ATAC-seq

Assay for Transposase-Accessible Chromatin by Sequencing (ATAC-seq) was used to profile open chromatin, as described previously with adaptations (Corces et al., 2017). XX_ΔXic_ and XO cells were profiled at day 0, 2 and 4 of differentiation in two biological replicates. Cells were dissociated with trypsin and 6*10^4^ cells were lysed in 50 μl cold RSB buffer (10 mM Tris-HCl (pH 7.4), 10 mM NaCl, 3 mM MgCl_2_, 0.1% Tween-20) supplemented with 0.1% Igepal CA-630 and 0.01% Digitonin. The lysis buffer was washed out using 1 ml of cold RSB buffer. Nuclei were then pelleted by centrifugation (500 x g, 10 min, 4 °C) and the supernatant aspirated. Subsequently, they were resuspended in 50 μl Transposase Mix (1x TD buffer (Illumina), 100 nM Nextera Tn5 Transposase (Illumina), 33 μl PBS, 0.01% Digitonin, 0.1% H_2_O) and incubated at 37°C for 30 minutes at 1000 rpm. The reaction was stopped by adding 2.5 μl 10% SDS and purified using the DNA Clean & Concentrator Kit (Zymo). 20 μl of the transposed DNA was then amplified using the NEBNext High-Fidelity 2x PCR Master mix with i5 and i7 Nextera barcoded primers for 12 cycles (see Supplementary Table 6 for primer sequences). The PCR product was size-selected using NGS Clean beads (CleanNA), by adding them at first at a 70%-ratio and transferring the supernatant. Afterwards, the beads were added once more at a 180%-ratio and the PCR product was eluted from the beads in 20 μl H_2_O. The success of the transposition was verified with the BioAnalyzer High Sensitivity DNA system (Agilent Technologies). Sequencing libraries were pooled in equimolar ratios and sequenced paired-end 75 bp on the HiSeq 4000 platform yielding approximately 2.5*10^7^ fragments per sample (Supplementary Table 1).

#### Data processing

Read sequences were trimmed using *Trim Galore* (v0.6.4) with options [--paired --nextera] (http://www.bioinformatics.babraham.ac.uk/projects/trim_galore/). Afterwards, the trimmed FASTQ files were aligned using *bowtie2* (Langmead and Salzberg, 2012) (v2.3.5.1) with the options [--local --very-sensitive-local -X 2000]. Mitochondrial reads were removed using a custom Python script. Mapping statistics can be found in Supplementary Table 1.

### STARR-seq

#### STARR-seq library cloning

A STARR-seq library covering the Xic was cloned as described previously (Arnold et al., 2013) with modifications. The Bacterial Artificial Chromosome (BAC) clones RP23-106C4, RP23-11P22, RP23-423B1, RP23-273N4, RP23-71K8 were purchased as bacterial stabs from the BAC PAC Resource Center of the Children’s Hospital Oakland Research Institute. *E.coli* BAC clones were grown in 200 ml LB medium (10 g/l NaCl, 10 g/l Bacto Tryptone, 5 g/l Yeast extract, 1 mM NaOH) supplemented with 12.5 μg/ml Chloramphenicol (Sigma) in a shaking incubator at 30 °C for 20 hours. The BAC DNA was isolated using the NucleoBond BAC 100 kit (Machery-Nagel). BAC DNA was pooled (2.5 μg each) and split into four tubes, which were filled with TE buffer to a total volume of 100 μl. The DNA was sheared by sonication (Bioruptor Plus, low intensity, 3 cycles with 32 sec ‘on’/ 28 sec ‘off’), size-selected on a 1% agarose gel and extracted with the QIAquick Gel Extraction Kit (Qiagen). The eluates were pooled and purified using the QIAquick PCR Purification Kit (Qiagen). The purified fragments were end-repaired, dA-tailed and ligated to adapter_STARR1/adapter_STARR2 to be compatible with Illumina sequencing according to the NEBnext DNA library prep master mix set for Illumina (New England Biolabs, oligonucleotide sequences shown in Supplementary Table 6). The ligated fragments were purified using Agencourt AMPure XP beads (Beckman Coulter) and eluted in 25 μl elution buffer. Four PCR reactions were then carried out with 1 μl of the purified DNA using the KAPA HotStart HiFi Ready Mix (KAPA Biosystems) for 10 cycles inserting a 15 nt homology sequence for the subsequent cloning step (IF_fwd/IF_rev). The PCR products were size-selected on a 1% agarose gel and purified using the MinElute PCR Purification Kit (Qiagen). The pSTARR-seq_human vector (kindly provided by Alexander Stark) was digested with AgeI-HF and SalI-HF for 3.5 h at 37 °C, size-selected on a 1% agarose gel and purified using the MinElute PCR Purification Kit (Qiagen). The DNA library was then cloned into the vector using four In-Fusion cloning reactions according to the manufacturer’s instructions (Clontech In-Fusion HD). The reactions were pooled, ethanol-precipitated and transformed into MegaX DH10BTM T1R Electrocompetent Cells (Invitrogen) according to the manufacturer’s instructions in a total of eight separate reactions. Plasmid DNA was amplified overnight and isolated using the NucleoBond Xtra Midi Plus Kit (Macherey-Nagel). The plasmid library was sequenced paired-end 50 bp on the HiSeq 2500 platform yielding approximately 1.1*10^7^ fragments (Supplementary Table 1). Due to a partial deletion in one of the BAC clones used, a ∼55 kb region within the *Linx* gene was not covered by the STARR-seq library.

#### Transfection and sequencing

5.0*10^6^ 1.8 XX and 1.8 XO cells were transfected with 2.5 μg of the STARR-seq library using Lipofectamine LTX (Thermo) according to manufacturer’s instructions in three biological replicates. 3*10^6^ cells were cultured under 2iL conditions and 2*10^6^ under differentiation conditions for 48 h. RNA was isolated using the Direct-Zol RNA Miniprep Kit (Zymo Research).The mRNA fraction was recovered from the total RNA using Dynabeads Oligo-dT25 (Invitrogen) with 1 mg beads per 50 μg of total RNA. The RNA was reverse transcribed into cDNA using Superscript III (Invitrogen) with a gene specific primer (STARR_GSP). Three reactions were performed for each sample. The reactions were then treated with RNaseI (Thermo) for 60 min at 37°C and cleaned using the MinElute PCR Purification Kit (Qiagen). Subsequently, a junction PCR was performed using an intron-spanning primer pair (Junction_fwd and Junction_rev) and 8 μl cDNA with the KAPA HotStart HiFi Ready Mix (KAPA Biosystems) for 15 cycles. Three reactions each were pooled using the QIAquick PCR Purification Kit (Qiagen). Sequencing adapters were added in a second PCR using three reactions with 10 μl of the purified junction PCR product with the KAPA HotStart HiFi Ready Mix (KAPA Biosystems) for 12 cycles. Lastly, the samples were isolated via agarose gel extraction and the QIAquick Gel Extraction Kit (Qiagen) and purified once more using the QIAquick PCR Purification and MinElute PCR Purification Kits (Qiagen). Libraries were pooled in equimolar ratios and sequenced paired-end 50 bp on the HiSeq 2500 platform yielding approximately 1.0*10^7^ fragments per sample (Supplementary Table 1). All primer sequences are provided in Supplementary Table 6.

#### STARR-seq data processing

The data was processed as described previously (Arnold et al., 2013). FASTQ files were mapped using *bowtie* (Langmead et al., 2009) (v1.2.2) with options [-S -t -v 3 -m 1 -I 250 -X 2000]. As the amount of reads per sample was very low after deduplication (∼99% duplicates) and the samples were similar between conditions, all samples were then merged using *samtools* (Li et al., 2009) (v1.10) with options [merge] for further analysis. For visualization, BIGWIG tracks normalized to the cloned library were created using *deepTools2* (Ramírez et al., 2016) (v3.4.1) with [bamCompare -e -bs 10 --operation ratio --normalizeUsing CPM]. Mapping statistics and quality control metrics are shown in Supplementary Table 1.

### CRISPRi screen

#### sgRNA library design

BAM files of all conditions and replicates of the ATAC-seq and STARR-seq data (see above) were merged. Peaks were called using *MACS2* (v2.1.2) with options [callpeak -f BAMPE -g mm -q 0.1] (Quinlan and Hall, 2010; Zhang et al., 2008). The resulting narrowPeak files were then filtered for peaks in the *Xic* (chrX:103198658-104058961). In addition, a list of candidate enhancer elements across different mouse tissues in the region, identified by the FANTOM5 consortium based on Cap Analysis of Gene Expression (CAGE), was used (Lizio et al., 2015). Afterwards, candidate regions from ATAC-seq, STARR-seq and FANTOM5 data were combined using *bedtools* (v2.29.2) with option [merge]. Regions longer than 2000 bp were split manually according to visual inspection of the ATAC-seq data and adjacent REs with a total combined length below 2000 bp (including the distance between them) were merged, resulting in a list of 138 candidate REs. Since the efficiency of targeting REs with CRISPRi is known to be highly variable (Klann et al., 2017), the candidate REs were saturated with sgRNA sequences generated from the *GuideScan* webtool (Perez et al., 2017) with a specificity score of >0.2 (Tycko et al., 2019). 300 randomly chosen non-targeting guides from the mouse CRISPR Brie lentiviral pooled library (Doench et al., 2016) were included as negative controls, resulting in 7358 guides in total.

#### sgRNA library cloning

The sgRNA library was cloned into the lentiGuide-puro sgRNA expression plasmid (Addgene 52963, (Sanjana et al., 2014)). The vector was digested with BsmBI (New England Biolabs) at 55 °C for 1 h and gel-purified. sgRNA sequences were synthesized by Genscript flanked with OligoL (TGGAAAGGACGAAACACCG) and OligoR (GTTTTAGAGCTAGAAATAGCAAGTTAAAATAAGGC) sequences. For the amplification of the library, 7 PCR reactions with primers OG113/OG114 and approx. 5ng of the synthesized oligo pool were carried out using the Phusion Hot Start Flex DNA Polymerase (New England Biolabs), with a total of 14 cycles and an annealing temperature of 63 °C in the first 3 cycles and 72 °C in the subsequent 11 cycles. The amplicons were subsequently gel-purified.

Amplified sgRNAs were ligated into the vector through Gibson assembly (New England Biolabs). Three 20 µl Gibson reactions were carried out using 7 ng of the gel-purified insert and 100 ng of the vector. The reactions were pooled, EtOH-precipitated to remove excess salts which might impair bacterial transformation and resuspended in 12.5 µl H_2_O. 9 µl of the eluted DNA were transformed into 20 µl of electrocompetent cells (MegaX DH10B, Thermo Fisher Scientific) according to the manufacturer’s protocol using the ECM 399 electroporator (BTX). After a short incubation period (1h, 37 °C 250 rpm) in 1 ml SOC medium, 9 ml of LB medium with Ampicillin (0.1 mg/ml, Sigma) were added to the mixture and dilutions were plated in Agar plates (1:100, 1:1000 and 1:10000) to determine the coverage of the sgRNA library (526-fold). 500 ml of LB media with Ampicillin were inoculated with the rest of the mixture and incubated overnight for subsequent plasmid purification using the NucleoBond Xtra Maxi Plus kit (Macherey & Nagel) following the manufacturer’s instructions. To assess library composition by deep-sequencing, a PCR reaction was carried out to add illumina adaptors by using the KAPA HiFi HotStart ReadyMix (Roche), with an annealing temperature of 60 °C and 12 cycles (OG125/OG126). The PCR amplicon was gel-purified by using the Nucleospin Gel and PCR Clean-up kit (Macherey & Nagel) following the manufacturer’s instructions. The library was sequenced paired-end 75 bp on the HiSeq 4000 Platform yielding approximately 7.5 million. fragments. Read alignment statistics found in Supplementary Table 2). A log2-distribution width of 1.7 for the plasmid library showed that sufficient coverage was attained during library cloning (Supplementary Fig. 1e, j). Only one sgRNA was missing from the library (gRNA_6494). All primer sequences are given in Supplementary Table 6.

#### Lentiviral packaging

HEK293T cells were cultured in DMEM supplemented with 10% FBS and passaged every 2 to 3 days. For lentiviral packaging, 20 10cm plates with HEK293T cells were transfected at 90% confluency, each with 6.3 µg pPL1, 3.1 µg pLP2 and 2.1 µg VSVG vectors (Thermo Fisher Scientific) together with 10.5 µg of the cloned sgRNA library. Plasmids and 60 µl Lipofectamine 2000 reagent (Thermo Fisher Scientific) were each diluted in 1 ml of OptiMEM, incubated separately for 5 min and then together for 20 min. The mix was added dropwise to the HEK293T cells and the medium was changed 6 h after transfection. After 48 h the medium was collected and viral supernatant was concentrated 10-fold using the lenti-X^TM^ Concentrator (Takara Bio) following the manufacturer’s instructions and subsequently stored at −80 °C.

To estimate the viral titer, serial 10-fold dilutions were prepared from the viral stock and used to transduce mESCs in a 6-well plate (Mock plus 10^-2^ to 10^-6^) together with 8 ng/µl polybrene (Merck) in duplicates. Selection with puromycin (1 ng/µl, Sigma) was started two days after transduction and colonies were counted in each well after 8 days. The estimated titer was 0.68*10^5^ transducing units (TU) per ml.

#### Lentiviral transduction

The TX-SP107 mESC line, carrying an ABA-inducible dCas9-KRAB system, was grown for at least two passages in SL medium prior to transduction. Transduction was carried out in SL medium, as X-chromosome loss was sometimes observed upon transduction in 2iL medium. A total of 6*10^6^ cells were transduced with viral supernatant of the sgRNA library (MOI = 0.3). Additionally, 2*10^5^ cells each were transduced with either an empty pLenti vector or an sgRNA targeting the *Xist* TSS (Supplementary Table 6). Both controls were taken along for the rest of the experiment and confirmed CRISPRi efficiency (Supplementary Fig. 1f-g). Puromycin selection (1 ng/µl, Sigma) was started two days after transduction and kept for the rest of the experiment. At the next passage, the cells were transferred into 2iL medium. After two more passages, cells were differentiated by 2iL-withdrawal. Recruitment of dCas9-KRAB to target sites was induced using ABA (100 µM) one day before differentiation and kept throughout the rest of the protocol. 1*10^6^ cells were kept in 2iL-containing medium and used as an undifferentiated control. Cells were harvested for Flow-FISH after 2 days of differentiation.

#### Flow-FISH and cell sorting

2*10^8^ cells were stained by Flow-FISH with an Xist-specific probe as described above. 2*10^7^ cells were snap-frozen after the two fixation steps to be used as the unsorted fraction. Four different populations were sorted, where 15% cells with the lowest signal were termed Xist-negative, while 45% cells with the strongest signal were sorted into 3 positive populations (0-15% = High, 15-30% = Medium, 30-45% = Low). Around 1.1-1.5*10^7^ cells were recovered per fraction. After sorting, the cell pellets were snap-frozen and stored at −80°C for further analysis.

#### Preparation of sequencing libraries and sequencing

Sequencing libraries were prepared from all sorted cell populations and the unsorted cells for each of the two independent screen replicates. DNA from frozen cell pellets was isolated through phenol/chloroform extraction since it yields significantly more DNA than DNA isolation kits based on silica columns. Cell pellets were thawed and resuspended in 250 µl of lysis buffer (1% SDS (Thermo Fisher Scientific), 0.2 M NaCl and 5 mM DTT (Roth) in TE Buffer) and incubated overnight at 65 °C. The next day 200 µg of RNase A (Thermo Fisher Scientific) were added and the samples were incubated at 37°C for 1 h. 100 µg of Proteinase K (Sigma) were subsequently added, followed by a 1 h incubation at 50°C. Phenol/chloroform/isoamyl alcohol (Roth) was added to each sample in a 1:1 ratio, the mixture was vortexed for 1 min and subsequently centrifuged at 16,000 x g for 10 min at RT. The aqueous phase was transferred to a new tube, 1 ml 100% EtOH, 90 µl 5 M NaCl and 1 µl Pellet Paint (Merck) was added to each sample, mixed, and incubated at −80 °C for 1 h. DNA was pelleted by centrifugation for 16,000 x g for 15 min at 4 °C, pellets were washed twice with 70% EtOH, air-dried and resuspended in 50 µl water.

The genomically integrated sgRNA cassette was amplified in two successive PCR reactions as described previously(Shalem et al., 2014) with minor modifications. To ensure sufficient library coverage (>300x), 14.5 µg of each sample were amplified using the ReadyMix Kapa polymerase (Roche) with a total of 20 cycles and an annealing temperature of 55 °C (OG115/OG116). Between 0.1-2 µg genomic DNA was amplified per 50 µl PCR reaction. In particular, in samples stained with Flow-FISH PCR amplification was inhibited at higher DNA concentrations such that up to 145 PCR reactions had to be performed per sample. Successful amplification was verified on a 1% agarose gel and the reactions were pooled. The PCR product was isolated and concentrated using the Zymo DNA Clean and Concentrator Kit. A second nested PCR was performed to attach sequencing adaptors and sample barcodes using 2.5-50 µl of the product from the first PCR as template, with a total of 11 cycles and an annealing temperature of 55 °C (OG125/OG170-OG180). Resulting amplicons were loaded on a 1% agarose gel and purified using the Nucleospin Gel and PCR clean-up kit (Macherey-Nagel). Libraries were sequenced paired-end 75bp on the NextSeq 500 platform yielding approximately 4*10^6^ fragments per sample (Supplementary Table 2). All primer sequences are given in Supplementary Table 6.

#### CRISPRi screen analysis

Data processing and statistical analysis was performed using the *MAGeCK* CRISPR screen analysis tools (Li et al., 2014, 2015) (v0.5.9.3). Alignment and read counting was performed with options [count --norm-method control] for all samples together. At least 3.25*10^6^ mapped reads were obtained per sample. Correlation between the two replicates was computed as a Pearson correlation coefficient on the normalized counts (Supplementary Fig. 1h). The NTC distribution width was similar across samples, suggesting that sufficient library coverage was maintained during all steps (Supplementary Fig. 1i-j).

Statistical analysis was performed on the RE levels with options [mle --norm-method control --max-sgrnapergene-permutation 350] and on the sgRNA level [test --norm-method control]. In order to rank REs based on their effect on *Xist* expression, we averaged their beta score, a measure of the effect size estimated by the *MAGeCK mle* tool, across populations for each RE that exhibited an FDR <0.05 in at least one bin, with inverting the sign in the negative bin. To ensure robustness of the ranking and to exclude an analysis bias associated with the variable number of sgRNAs per RE, we implemented an alternative strategy focussing on those REs that were targeted by >50 guides. First, normalized counts were averaged across replicates for each sgRNA. For 1000 bootstrap samples, each containing 50 sgRNAs randomly selected with replacement, the fold change between sorted and unsorted fractions was calculated and averaged. Ranking REs according to the mean of those fold-change distributions led to nearly identical results as the beta-score based approach. An empirical p-value was calculated from the resulting distribution and Benjamini-Hochberg corrected. Alignment statistics, normalized counts, gene hit summary files and RE ranking is provided in Supplementary Table 2.

### CUT&Tag of histone modifications

Cleavage Under Targets and Tagmentation (CUT&Tag) makes use of Tn5 transposition at protein A (pA) bound antibody recognition sites and was performed as described previously with minor modifications (Kaya-Okur et al., 2019).

#### Purification of 3xFLAG-pA-Tn5

For this we purified 3xFlag-pA-Tn5 from E. coli containing pTXB1-3xFlag-pA-Tn5-FL (Addgene, #124601), a kind gift from (Kaya-Okur et al., 2019). From an overnight streak LB agar plate, a single colony was selected for a liquid starter culture in LB medium supplemented with Carbenicilin (100 μg/ml) and incubated in a shaker at 37 °C for four hours. Afterwards the starter culture was added to 400 ml LB medium supplemented with Carbenicilin (100 μg/ml) and incubated until it reached an OD_600_ of 0.6 (roughly three hours) and was directly cooled on ice. After 30 min on ice, 100 μl of 1 M IPTG was added to the culture and incubated in a cooled shaker overnight at 18 °C at 150 rpm. The following morning, bacteria were centrifuged in a JA-12 rotor at 10,000 rpm for 30 min at 4 °C. Bacterial pellets were snap frozen in liquid nitrogen and stored at −80 °C. The pellets were thawed on ice and resuspended in 40 ml HEGX buffer (20 mM HEPES-KOH pH 7.2, 0.8 M NaCl, 1 mM EDTA pH8.0, 10% Glycerol, 0.2% Triton X-100). Following this the cell suspension was divided over two 50 ml tubes and lysed with a Branson tip sonicator on ice by using a 1064 (10-150 ml) tip with the following settings: 20 sec on, 20 sec off, 50% duty cycle for 9 min total. Afterwards the lysate was centrifuged in a JA-12 rotor at 10,000 rpm for 30 min at 4 °C and the supernatant was collected. Two 20 ml columns (Biorad, #7321010) were each packed with 2.5 ml Chitin resin slurry (New England BioLabs) and washed once with 20ml HEGX buffer. To each column 20 ml supernatant was added, locked on both openings and incubated overnight with rotation. The following morning the columns were washed four times with 20ml pre-cooled HEGX buffer supplemented with protease inhibitor cocktail tablets (Roche). Afterwards the chitin resin holding the 3xFLAG-pA-Tn5 was collected in a total of 10 ml elution buffer (20 mM HEPES-KOH pH7.2, 0.8 M NaCl, 1 mM EDTA, 10% Glycerol, 0.2% Triton X-100, 100 mM DTT), transferred to a 15ml falcon tube and extracted for 48 h on a rotator (15 rpm) at 4 °C. Afterwards the resin was allowed to settle to the bottom over 40 min on ice followed by centrifugation for two min at 300 rpm at 4 °C to collect all chitin resin. The supernatant holding the 3xFLAG-pA-Tn5 was dialysed in 800 ml cold dialysis buffer (100 mM HEPES-KOH pH7.2, 0.1 M NaCl, 0.2 mM EDTA, 2 mM DTT, 0.2% Triton X-100, 20% Glycerol) using a Slide-A-lyzer 30K dialysis cassette for 24 h at 4 °C with magnetic stirring at 300 rpm, with the buffer being refreshed after the initial 12 h. The dialysed protein extract (∼5.5ml) was concentrated 6-fold using a Amicon Ultra 4 30K 15ml falcon filtration system with successive rounds of centrifugation in a swing bucket centrifuge at 3000 xg at 4 °C for 15 min. The protein concentration was measured with the detergent compatible Bradford assay kit (Pierce), adjusted to 832 ng/μl with dialysis buffer and diluted 1:1 volume with 100% glycerol (= 5.5 µM). The 3xFLAG-pA-Tn5 fusion protein was confirmed on a GelCode Blue (Thermo Fisher Scientific) stained SDS-PAGE. Aliquots of 100 μl of 5.5 µM 3xFLAG-pA-Tn5 fusion protein were loaded with mosaic end adapters. For this, 10 μl ME-A with 10 μl ME-reverse and 10 μl ME-B with 10μl ME-reverse 200 µM oligos (Supplementary Table 6) (dissolved in 10 mM Tris-HCl pH8.0, 50 mM NaCl, 1 mM EDTA) were annealed in separate reactions on a thermocycler for 5 minutes at 95 °C with a ramp down to 21 °C over 30 min and mixed together afterwards. Aliquots of 100 μl containing 5.5 µM 3xFLAG-pA-Tn5 fusion protein were mixed with 16 µl of adapter mix and incubated for 1 h at RT with rotation and stored at −20 °C.

#### Cell preparations and CUT&Tag

Cells were washed with PBS and dissociated with accutase. For each CUT&Tag reaction 1*10^5^ cells were collected and washed once with Wash buffer (20 mM HEPES-KOH, pH 7.5, 150 mM NaCl, 0.5 mM Spermidine, 10 mM Sodium butyrate, 1 mM PMSF). 10 μl Concanavalin A (Bangs Laboratories) beads were equilibrated with 100 μl Binding buffer (20 mM HEPES-KOH, pH 7.5, 10 mM KCl, 1 mM CaCl_2_, 1 mM MnCl_2_) and afterwards concentrated in 10 μl binding buffer. The cells were bound to the Concanavalin A beads by incubating for 10 min at RT with rotation. Following this, the beads were separated on a magnet and resuspended in 100 μl chilled Antibody buffer (Wash buffer with 0.05% Digitonin and 2 mM EDTA). Subsequently 1 µl of primary antibody (antibodies can be found in Supplementary Table 6) was added and incubated on a rotator for 3 hours at 4 °C. After magnetic separation the beads were resuspended in 100 μl chilled Dig-wash buffer (Wash buffer with 0.05% Digitonin) containing 1 µl of matching secondary antibody and were incubated for 1 h at 4 °C with rotation. The beads were washed three times with ice cold Dig-wash buffer and resuspended in chilled Dig-300 buffer (20 mM HEPES-KOH, pH 7.5, 300 mM NaCl, 0.5 mM Spermidine, 0.01% Digitonin, 10 mM Sodium butyrate, 1 mM PMSF) with 1:250 diluted 3xFLAG-pA-Tn5 preloaded with Mosaic-end adapters. After incubation for 1 h at 4 °C with rotation, the beads were washed four times with chilled Dig-300 buffer and resuspended in 50 μl Tagmentation buffer (Dig-300 buffer 10 Mm MgCl_2_). Tagmentation was performed for 1 h at 37 °C and subsequently stopped by adding 2.25 µL 0.5 M EDTA, 2.75 µL 10% SDS and 0.5 µL 20 mg/mL Proteinase K and vortexing for 5 sec. DNA fragments were solubilized overnight at 55 °C followed by 30 min at 70 °C to inactivate residual Proteinase K. DNA fragments were purified with the ChIP DNA Clean & Concentrator kit (Zymo Research) and eluted with 25 μl elution buffer according to the manufacturer’s guidelines.

#### Library preparation and sequencing

NGS libraries were generated by amplifying the CUT&Tag DNA fragments with i5 and i7 barcoded HPLC-grade primers (Buenrostro et al., 2015) (Supplementary Table 6) with NEBNext® HiFi 2x PCR Master Mix (New England BioLabs) on a thermocycler with the following program: 72 °C for 5 min, 98 °C for 30 sec, 98 °C for 10 sec, 63 °C for 10 sec (14-15 Cycles for step 3-4) and 72 °C for 1 min. Post PCR cleanup was performed with Ampure XP beads (Beckman Coulter). For this 0.95x volume of Ampure XP beads were mixed with the NGS libraries and incubated at RT for 10 min. After magnetic separation, the beads were washed three times on the magnet with 80% ethanol and the libraries were eluted with Tris-HCl, pH 8.0. The quality of the purified NGS libraries was assessed with the BioAnalyzer High Sensitivity DNA system (Agilent Technologies). Sequencing libraries were pooled in equimolar ratios, cleaned again with 1.2x volume of Ampure XP beads and eluted in 20 μl Tris-HCl, pH 8.0. The sequencing library pool quality was assessed with the BioAnalyzer High Sensitivity DNA system (Agilent Technologies) and subjected to Illumina PE75 next generation sequencing on the NextSeq500 platform totalling approximately 5 million fragments per library.

#### Data processing

Read sequences were trimmed using *Trim Galore* (0.6.4) with options [--paired --nextera] (http://www.bioinformatics.babraham.ac.uk/projects/trim_galore/). Afterwards, the trimmed FASTQ files were aligned according to (Kaya-Okur et al., 2019) with modifications using *bowtie2* (v2.3.5.1) with the options [--local --very-sensitive-local --no-mixed --no-discordant --phred33 -I 10 -X 2000] (Langmead and Salzberg, 2012). As the percentage of duplicate reads was very low, duplicated reads were kept for analysis. Mapping statistics and quality control metrics can be found in Supplementary Table 3.

#### Correlation analysis

BAM files, excluding mitochondrial reads, were counted in 1 kb bins using *deepTools2* (Ramírez et al., 2016) (v3.4.1) with options [multiBamSummary bins -bs 1000 -bl chrM.bed]. The Pearson correlation coefficient between different histone marks, conditions or replicates was then computed using *deepTools2* (v3.4.1) with options [plotCorrelation -c pearson]. The resulting values were hierarchically clustered and plotted as a heatmap.

#### Genomic peak annotation

Location of CUT&Tag peaks was analyzed using *ChIPseeker* (Yu et al., 2015) (v1.22.1) in undifferentiated XX_ΔXic_ mESCs. Peaks identified using *MACS2* (see above) were assigned to gene features according to the annotation package *TxDb.Mmusculus.UCSC.mm10.knownGene* (v3.10.0) (Supplementary Fig. 1f) (https://bioconductor.org/packages/TxDb.Mmusculus.UCSC.mm10.knownGene/).

#### Comparison of CUT&Tag with native ChIP-seq data

The H3K4me1, H3K4me3, H3K27ac, H3K27me3 and H2AK119ub histone marks profiled via CUT&Tag in undifferentiated XX_ΔXic_ cells were compared to native ChIP-seq data profiling the same marks in the parental TX1072 cell line (Żylicz et al., 2019). FASTQ files were retrieved from the GEO Accession Viewer (GSE116990) using *fasterq-dump* (v2.9.4) (http://ncbi.github.io/sra-tools/). In order to keep the data comparable, processing was done analogous to the CUT&Tag data, as described above. Subsequently, reads were quantified in 1 kb bins using *deepTools2* (Ramírez et al., 2016) (v3.4.1) with the options [multiBamSummary -bs 1000] on merged replicates. Afterwards, a PCA analysis was conducted using the base R package *stats* (v3.6.3) with [prcomp(center = TRUE, scale = TRUE)].

### Differential peak analysis

Differential peaks were identified for ATAC-seq, H3K4me3, H3K4me1 and H3K27ac with the *DiffBind* bioconductor package (Ross-Innes et al., 2012) (v2.6.6). The analysis was performed either for all peaks that were identified with *MACS2* (see above) in all replicates of at least one condition or for all candidate REs from the CRISPR screen (Fig. 3d). All peaks on the X chromosome outside of the deleted region in the XX_ΔXic_ cell line (chrX: 103182701-103955531) were excluded from the analysis to remove any potential bias due to the different number of X chromosomes between the cell lines. Differential peaks were then analyzed between timepoints and cell lines with the options [dba.analyze(method = DBA_ALL_METHODS)]. The results were exported using [dba.report(method = DBA_ALL_METHODS, th = 0.05, bUsepVal = FALSE)].

In order to find differentially enriched regions for the broader H3K9me3 mark, we used *diffReps* (Shen et al., 2013) (v1.55.6). Here the number of reads mapping to 5 kb intervals was compared between time points or cell lines using a sliding window approach with options [--window 5000 --step 1000]. To identify consensus peaks on the X chromosome present in both conditions, *diffreps* was used to compare each condition with a modified version of itself, where all X-chromosomal reads had been removed. Peaks identified in both conditions, which did not overlap with a differential peak were defined as consensus peaks..

### ChromHMM analysis

Chromatin segmentation was performed using *ChromHMM* (Ernst and Kellis, 2012) (v1.19) on ATAC-seq and CUT&Tag data (H3Kme1, H3Kme3, H3K27ac, H3K27me3) for the XX_ΔXic_ cell line at all three timepoints. The model was learned for 10 to 15 emission states. After visual inspection of the resulting BED files, 12 emission states were chosen for further analysis. Chromatin states were then assigned as ‘no RE’, ‘poised RE’, ‘weak RE’ or ‘strong RE’ states depending on the enrichment of the different chromatin marks (Supplementary Fig. 2h).

### Quantification of sequencing data in candidate REs

ATAC-seq, H3K4me3, H3K27ac and H3K4me1 reads were quantified from the replicate BAM files at the candidate REs using *Rsubread* (Liao et al., 2019) (v2.0.1) with the options [featureCounts(isPairedEnd = TRUE)]. Counts per Million (CPM) were then calculated for all samples. To compare between different conditions, we computed a z-score for the individual REs (Fig. 3c). In Fig. 3c,d, comparisons in which all of the conditions failed to reach 5 raw reads in both replicates were colored in dark gray.

### Published ChIP-seq data

FASTQ files for transcription factor binding data of SMAD2/3 and TCF3 in embryoid bodies and ESCs (Wang et al., 2017), of OCT4 and OTX2 in Epi-like stem cells (EpiLC) and ESCs (Buecker et al., 2014) and of CTCF in ESCs (Stadler et al., 2011) were retrieved from the GEO Accession Viewer (GSE70486, GSE56098 and GSE30203) using *fasterq-dump* (v2.9.4) (http://ncbi.github.io/sra-tools/). Processed WIG tracks of ChIP-seq data for H3K4me1, H3K4me3 and H3K27ac at timepoints E6.5, E7.0 and E7.5 (Yang et al., 2019) were retrieved from the GEO Accession viewer (GSE98101). Reads were trimmed for adapter fragments using *Trim Galore* (v0.6.4) with options [--illumina] (http://www.bioinformatics.babraham.ac.uk/projects/trim_galore/). Reads were aligned using *bowtie2* (Langmead and Salzberg, 2012) (v2.3.5.1) with the options [--local --very-sensitive-local --no-mixed --no-discordant --phred33 -I 10 -X 2000] for paired-end and [--very-sensitive] for single-end data (Langmead and Salzberg, 2012)

#### Visualization of CTCF binding sites within the Xic

CTCF binding sites (CBS’s) in mESC’s were visualized using the *FIMO* program within the *MEME suite* web tool (Grant et al., 2011) (v5.2.0). To this end, we generated a FASTA file containing the sequences within the CTCF peaks using *bedtools* (Quinlan and Hall, 2010) (v2.29.2) with options [getfasta]. Then we scanned the peaks for the occurence of the CTCF transcription factor binding motif, which was retrieved from the *JASPAR* database (Fornes et al., 2020) (8^th^ release). Lastly, the direction of the CBS’s were annotated by the strandedness of the binding motif.

### TT-seq and RNA-seq

#### S4U metabolic labeling of nascent RNA

Transient transcriptome sequencing (TT-seq), which is based on enrichment of S4U-labeled nascent RNA after a short, 5 minute labelling pulse (Schwalb et al., 2016), was performed to profile genome-wide nascent RNA levels. To this end the cells were cultured in 10 cm plates with 2iL or differentiated (2iL withdrawal) for 2 or 4 days. Cells were metabolically labeled with culture medium containing 750 μM 4-Thiouridine (S4U) (Sigma Aldrich) for 5 min at 37 °C and 5% CO_2_. Directly afterwards, the cells were washed with PBS and lysed on ice with TRIzol (Ambion).

#### RNA isolation

The lysates were pre-cleared by centrifugation and per 1*10^7^ cells supplemented with 2.4 ng equimolar mix of *in vitro* transcribed S4U-labelled and unlabeled ERCC spike-ins, as previously described (Schwalb et al., 2016). Total RNA was extracted from TRIzol with chloroform. In short, 200 μl chloroform was mixed per ml lysate and phase separated by centrifugation in phase-lock tubes. RNA was precipitated from the aqueous phase with isopropanol supplemented with 0.1 mM DTT and centrifugation for 10 min at 16.000x g and 4 °C. The RNA pellet was washed once with 75% ethanol and resuspended in nuclease free water. Residual genomic DNA was removed in solution with DNaseI (Qiagen) following the manufacturer’s guidelines. The total RNA was purified for a second round with the Direct-zol RNA Miniprep Plus kit (Zymo Research) by following the manufacturer’s guidelines but in addition including 100 mM DTT in all wash buffers to prevent oxidation of S4U-labeled RNA.

#### RNA fragmentation and biotinylation

For each TT-seq reaction 300 μg of purified RNA was divided over 2 Covaris MicroTubes and fragmented on the Covaris S2 platform for 10 s, 1% duty cycle, intensity 2, 1 cycle, 200 cycles/burst. Corresponding samples were pooled and 3 μg taken aside for quality control (detailed below). The remaining S4U-treated RNA (∼260 μl) was biotinylated by adding 240 μl nuclease free water, 100 μl 10x Biotinylation buffer (100 mM Tris-HCl pH7.4, 10 mM EDTA), 200 μl DMSO and 200 μl Biotin-HPDP (1 μg/μl in DMSO) and incubated for 1.5 h on a thermoblock at 37 °C with 750 rpm agitation. To remove unreacted Biotin-HPDP, the biotinylated RNA was extracted with phenol:chloroform (PCI) 5:1, pH 4.5) (Ambion). For this an equal volume of PCI was mixed with the biotinylated RNA and phase separated by centrifuging in phase-lock tubes at 12,000 x *g* for 5 min. The aqueous phase was collected, mixed with an equal volume of isopropanol and 1:10 volume of 5 M NaCl, and centrifuged at 16,000 x *g* for 15 min 4 °C to precipitate the biotinylated RNA. The RNA pellet was washed twice with 500 μl 75% ethanol and dissolved in 50 μl RNase-free water.

#### Nascent RNA enrichment

Biotinylated RNA was denatured at 65 °C for 10 min, directly followed by cooling on ice. To capture the biotinylated (nascent) RNA, 100 μl μMACS Streptavidin Microbeads (Miltenyi) were added and incubated on a heat block at 24 °C for 15 min with 750 rpm agitation. Bead mixture was loaded on pre-equilibrated MACS μColumn while attached to a μMACS separator. The initial flow-through was collected and loaded one more time on the MACS μColumn. The columns were washed 3 times with 900 μl heated (65 °C) wash buffer (100 mM Tris-HCl pH 7.4, 10 mM EDTA, 100 mM NaCl, 0.1% Tween-20) and 3x with RT wash buffer. To elute the enriched nascent RNA, the columns were loaded twice with 100 μl 100 mM DTT and collected. The nascent RNA was purified by adding 3 volumes of TRIzol and processed with the Direct-zol RNA Miniprep kit (Zymo Research) with addition of 1/100 volume of 100 mM DTT to each supplied wash buffer.

To confirm the quality of the total input and nascent RNA, the samples were analyzed with the Agilent RNA 600 pico kit on the Bioanalyzer platform (Agilent). Furthermore, enrichment of labelled (nascent) RNA over unlabeled RNA was assessed by RT-qPCR. For this, 1 µl of eluted nascent RNA and 500 ng fragmented total RNA were reverse transcribed (as described above) and enrichment of labelled ERCC spike-ins over unlabelled spike-ins (included during the first RNA isolation steps) was assessed by qRT-PCR (Supplementary Fig. 4a) with primers specific for each spike-in sequence (Supplementary Table 6).

#### Library preparation and sequencing

Total RNA and nascent RNA samples were subjected to strand-specific RNA-seq library preparation with the KAPA RNA HyperPrep kit with RiboErase (Illumina), which included 1st and 2nd strand synthesis and ribosomal RNA depletion, by following the manufacturer’s guidelines. The libraries were sequenced PE75 (for XX_ΔXic_) on the Illumina HiSeq 4000 platform or PE100 (for XO) on the NovaSeq 6000 platform with approximately 25 million fragments for total RNA and 100 million fragments for nascent RNA.

#### Data processing

Total and nascent RNA data was processed according to Schwalb et al.(Schwalb et al., 2016). Reads were aligned using *STAR* (v2.7.5a) with options [--outSAMattributes NH HI NM MD] (Dobin et al., 2013). Mapping statistics and quality control can be found in Supplementary Table 4.

#### Gene quantification

To quantify gene expression the GENCODE M25 annotation(Frankish et al., 2019) was supplemented with the *Xert* coordinates (Supplementary Table 5). *Rsubread* (Liao et al., 2019) (v2.0.1) was used with the options [featureCounts(isPairedEnd = TRUE, GTF.featureType = “gene”, strandSpecific = 2, allowMultiOverlap = TRUE)] to count read over the entire gene for TT-seq or with [featureCounts(isPairedEnd = TRUE, GTF.featureType = “exon”, strandSpecific = 2, allowMultiOverlap = FALSE)] to only count exonic reads for RNA-seq. In order to detect statistical differences in the expression of lncRNA expression within the *Xic*, we performed differential expression analysis between the XX_ΔXic_ and XO cell lines using *DESeq2* (Love et al., 2014) (v1.26.0). Comparisons with an adjusted p-value <0.05 were marked as significant (Supplementary Fig. 4b). TPM values and the results of the differential expression analysis can be found in Supplementary Table 4.

RNA-seq (SMART-seq2) data for embryonic mouse tissues at different stages of development was retrieved from the GEO Accession Viewer (GSE76505) (Zhang et al., 2018) and analyzed in the same way.

### Capture Hi-C

#### Nuclei preparation

XX_ΔXic_ and XO cultured in 2iL (day 0) or after 2 days differentiation (2iL withdrawal) were disassociated with 0.1% (w/v) accutase for 7 min at 37 °C. Cells were counted and 2*10^6^ cells were transferred in a 50 ml falcon tube through a 40 μm cell strainer and complemented with 10% FBS in PBS. 37% formaldehyde (Sigma-Aldrich) was added to a final concentration of 2% to fix the cells for 10 min at RT. Crosslinking was quenched by adding glycine to a final concentration of 125 mM. Fixed cells were washed twice with cold PBS and lysed using fresh lysis buffer (10 mM Tris, pH 7.5, 10 mM NaCl, 5 mM MgCl_2_, 0.1 mM EGTA with protease inhibitor cocktail tablets (Roche) to isolate nuclei. After 10 min incubation in ice, cell lysis was assessed microscopically. Nuclei were centrifuged for 5 min at 480 x *g*, washed once with PBS and snap frozen in liquid N_2_.

#### Chromosome conformation capture library preparation and sequencing

3C libraries were prepared from fixed nuclei as described previously (Despang et al., 2019). In summary, nuclei pellets were thawed on ice and subjected to DpnII digestion, ligation and decrosslinking. Re-ligated products were sheared using a Covaris sonicator (duty cycle: 10%, intensity: 5, cycles per burst: 200, time: 2 cycles of 60 sec each, set mode: frequency sweeping, temperature: 4–7 °C). Adapters were then added to the sheared DNA and amplified according to Agilent instructions for Illumina sequencing. The library was hybridized to the custom-designed SureSelect library and indexed for sequencing (100 bp, paired end) following manufacturer’s instructions. The custom-designed SureSelect library was described to capture informative GATC fragments within chrX:102238718-105214261 (mm10) using *GOPHER*, as described previously(Hansen et al., 2019). Capture Hi-C experiments were performed in duplicate which displayed strong replicate correlation (Supplementary Fig. 7e).

#### Processing of cHi-C experiments

Mapping, filtering and deduplication of short reads were performed with the *HiCUP* pipeline (Wingett et al., 2015) (v0.7.4) [no size selection, Nofill: 1, Format: Sanger]. The pipeline employed *bowtie2* (v2.3.5.1) (Langmead and Salzberg, 2012) for mapping short reads to the N-masked reference genome mm10. *Juicer tools* (v1.19.02) (Durand et al., 2016) was used to generate binned contact maps from valid and unique read pairs with MAPQ ≥ 30 (Durand et al., 2016) and to normalize contact maps by Knight and Ruiz (KR) matrix balancing (Knight and Ruiz, 2013). For the generation of contact maps, only read pairs mapping to the genomic region chrX:103,190,001-103,950,000 were considered. In this part of the enriched region, both investigated cell lines (XO and XX_ΔXic_) have only one allele. Afterwards, KR-normalized maps were exported at 10 kb bin size.

Subtraction maps were generated from KR-normalized maps, which were normalized in a pairwise manner before subtraction. To account for differences between two maps in their distance-dependent signal decay, the maps were scaled jointly across their sub-diagonals. Therefore, the values of each sub-diagonal of one map were divided by the sum of this sub-diagonal and multiplied by the average of these sums from both maps. Afterwards, the maps were scaled by 10^6^/total sum.

cHi-C maps were visualized as heatmaps with values above the 0.92-quantile being truncated to improve visualisation. For subtraction maps, the 0.95-quantile was computed on absolute values and used as truncation threshold for positive values as well as for negative values (with minus sign).

## Data Availability

All RNA-seq, CUT&Tag, TT-seq, ATAC-seq, STARR-seq, Capture-HiC and CRISPR screen data sets generated during this study are available on GEO (Accession number: GSE167358). All code used to analyse the NGS data is available at Github (https://github.com/EddaSchulz/Xert_paper/).

## Acknowledgements

We want to thank Abhishek Sampath Kumar for help with generating the E14-STN_ΔTsixP_ line, Steven Henikoff for support with setting up CUT&Tag, Kristina Zumer for support with setting up TT-seq. We thank Tyler Klann and Charles Gersbach for support with setting up the CRISPR screen, Laura Glaser and Melissa Bothe for support with establishing ATAC-seq and Pablo Navarro for sharing the E14 SunTag cell line. We thank Rafael Galupa for sharing the pFX5 plasmid and critical reading of the manuscript. We thank the Max Planck Institute for Molecular Genetics Seqcore and FACS facilities. This work was supported by the Max-Planck Research Group Leader program, E:bio Module III—Xnet grant (BMBF 031L0072) and Human Frontiers Science Program (CDA-00064/2018) to E.G.S. T.S. is supported by the DFG (IRTG2403, Regulatory Genome). V.M. is supported by the DFG (GRK1772, Computational Systems Biology).

## Author contributions

RAFG, TS and EGS conceived the project and designed the experiments. RAFG performed CUT&Tag, TT-seq and CRISPRi/a experiments. TS performed ATAC-seq, CRISPRi screen and CRISPRi Flow-FISH. LRL established the Flow-FISH screening assay. LB performed STARR-seq. LRL and VS generated CRISPRa/i multi-guide plasmids. ID and VS generated CRISPRi/a mESC lines. VM generated the TX1072 XX_ΔXic-B6_ line with help from RAFG. PK, RAFG and TS generated and analyzed mESC mutant lines. RACE was performed by PK and *de novo* transcript assembly by EN. MR performed cHiC and RS analysed the data. CV performed differential binding analysis for CUT&Tag. TS performed all other computational analyses. RAFG, TS and EGS wrote the manuscript with input from all authors. Funding acquisition by AM, SM and EGS.

## Supplementary Material

**Supplementary Figure 1.**
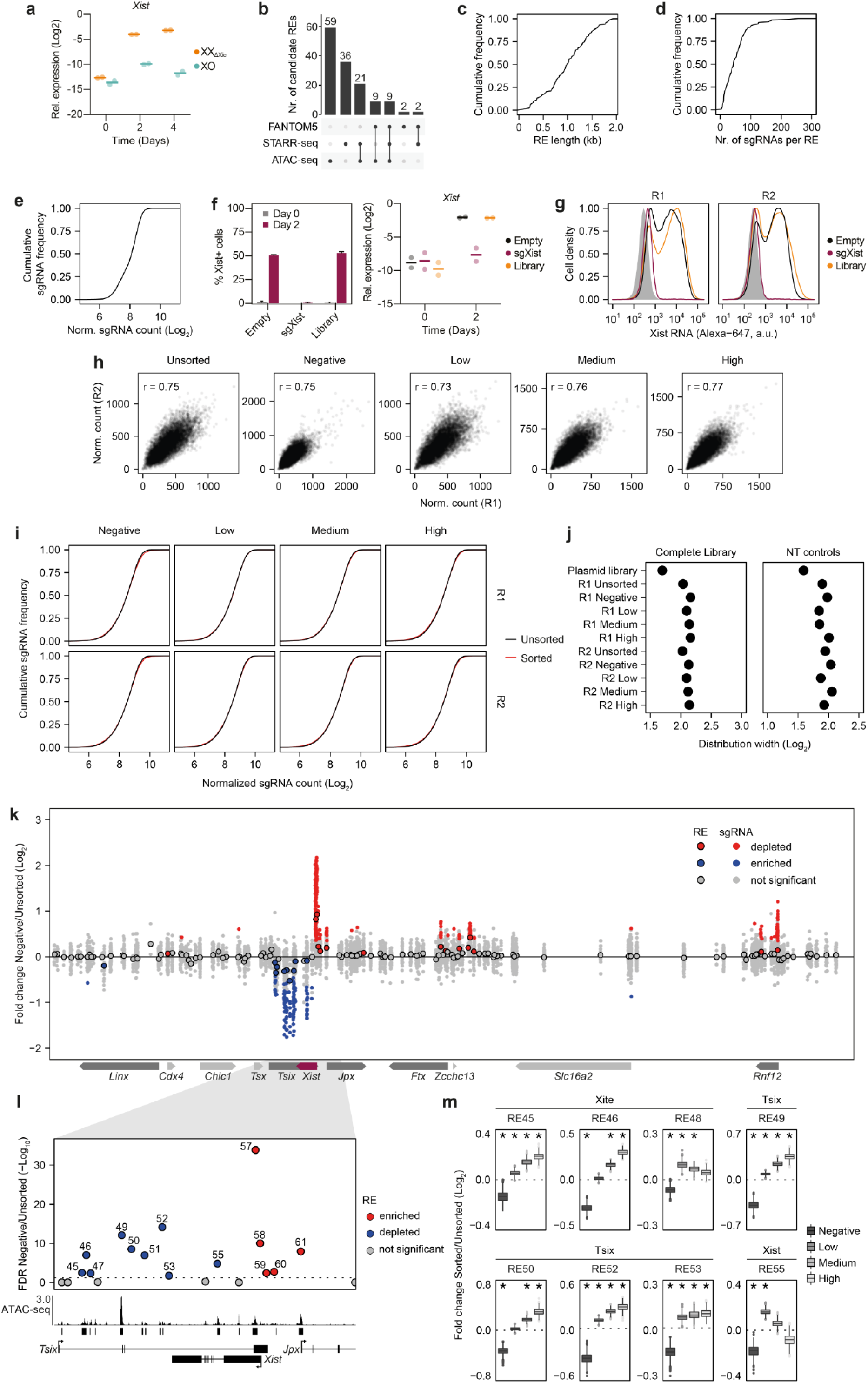
Identification of *Xist*-regulating genomic elements through a pooled CRISPR screen. (**a**) *Xist* expression assessed by RT-qPCR as quality control, when generating ATAC-seq data showing XX-specific upregulation during differentiation. Dots indicate individual biological replicates (n=2) and horizontal bars the mean. (**b**) UpsetR plot showing the number of candidate REs identified from the different data sources. (**c-d**) Cumulative frequency plots showing the distribution of length (c) and the number of sgRNAs (d) across all candidate REs. (**e**) Cumulative frequency plot showing the distribution of sgRNA counts in the cloned sgRNA library. (**f-g**) Quality controls for the CRISPR screen in Fig. 1 confirming *Xist* upregulation upon transduction with the sgRNA library (Library) or the empty vector (Empty) and Xist repression by a control sgRNA targeting the *Xist* promoter (sgXist). *Xist* was quantified in two biological replicates by RNA-FISH (f, left), RT-qPCR (f, right) and by Flow-FISH (g). For RNA-FISH, 100 cells were counted per replicate and mean and s.d. are shown. For Flow-FISH both replicates (R1, R2) are shown and undifferentiated cells transduced with the sgRNA library are shaded in grey. (**h**) Scatterplots showing the correlation between the replicates in the screen for each fraction as indicated. Pearson correlation coefficients are indicated. (**i**) Cumulative frequency plots showing the distribution of the sgRNA library in the sorted fractions compared to the unsorted population. (**j**) Log2 distribution width (fold change between the 10^th^ and 90^th^ percentiles) for all sgRNAs (left) and non-targeting (NT) sgRNAs only (right). The NT distribution width was similar across samples, suggesting that sufficient library coverage was maintained during all steps of the screen. (**k-l**) Comparison of sgRNA abundance in the Xist-negative fraction compared to the unsorted population. Small dots in (k) show individual sgRNAs and rimmed circles in (k-l) denote results from a joint analysis of all sgRNAs targeting one RE. Significantly enriched and depleted sgRNAs (*MAGeCK test,* two sided p-value <0.05) and REs (*MAGeCK mle,* Wald.FDR <0.05) are colored blue and red, respectively. The entire region targeted in the screen is shown in (k) and a zoom-in around Xist in (l). In (l) ATAC-seq data is shown from differentiated XX_ΔXic_ cells at day 2. (**m**) Log2 fold-change of sorted and unsorted populations for 1000 bootstrap samples of 50 randomly selected sgRNAs. REs in TAD-D with an empirical FDR <0.01 (asterisks) in at least two populations are shown.

**Supplementary Figure 2.**
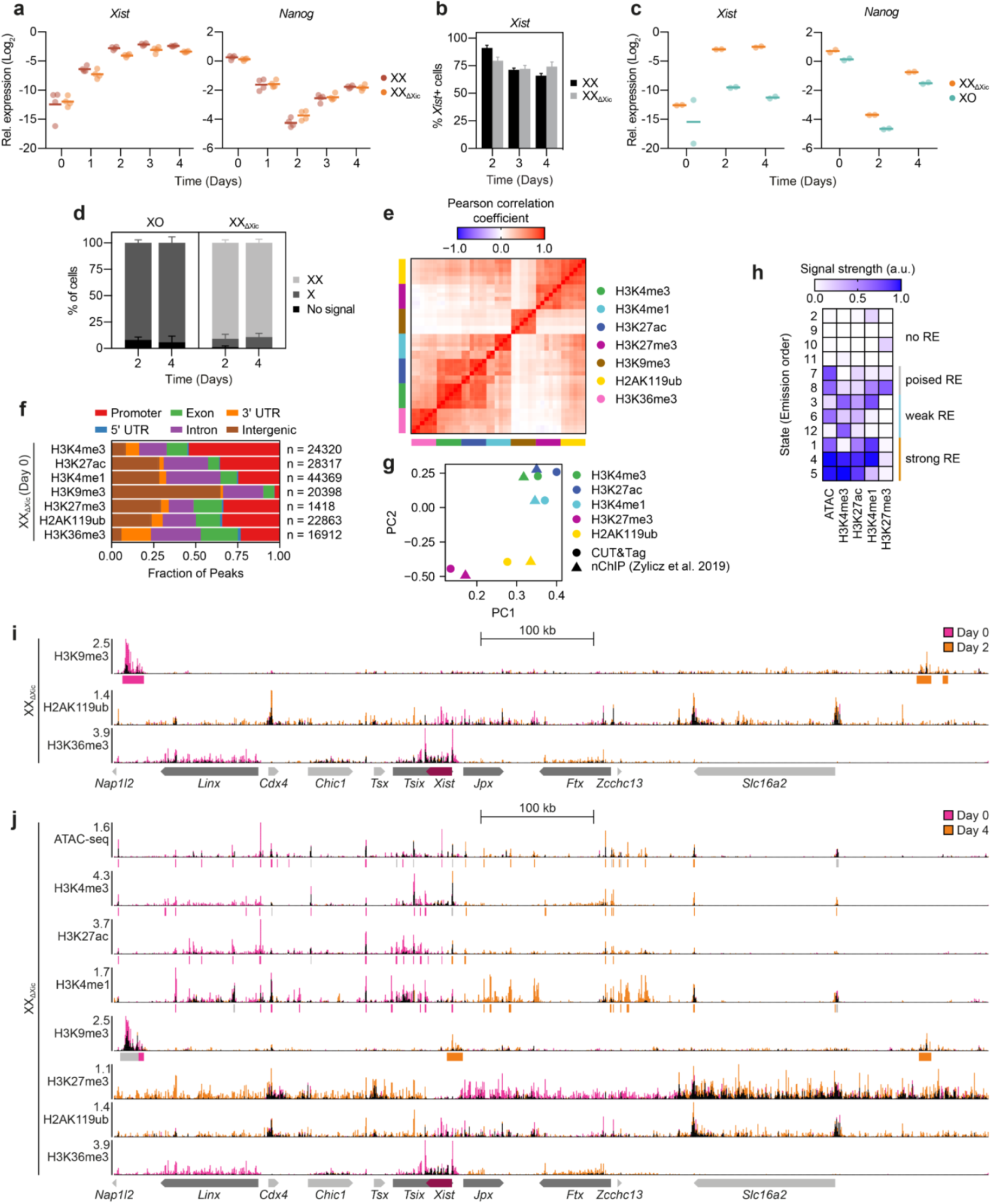
Differentiation cues affect distal, but not proximal *Xist*-controlling elements. (**a-b**) RT-qPCR and RNA-FISH comparing the XX_ΔXic_ line with the parental TX1072 cell line. In (a) dots represent 4 independent biological replicates, horizontal bars the mean. In (**b**) mean and s.d. of 3 biological replicates are shown. (**c-d**) Quality controls for the differentiation experiments, where chromatin was profiled by CUT&Tag. (c) RT-qPCR showing expression of *Xist* and *Nanog*. Dots represent 2 independent biological replicates, horizontal bars the mean. (**d**) Number of X chromosomes quantified by RNA-FISH for an X-linked gene (*Huwe1*). 100 cells were counted per replicate. Mean and s.d. of 2 biological replicates are shown. (**e**) Heatmap showing the Pearson correlation coefficients between all CUT&Tag samples. Replicates were merged and samples are ordered via hierarchical clustering. The expected correlation pattern was observed among modifications associated with active genes (H4K4me1/3, H3K27ac, H3K36me3) and among those associated with polycomb-repression (H3K27me3, H2AK119ub). (**f**) Distribution of CUT&Tag peaks in undifferentiated XX_ΔXic_ mESCs across genomic regions as indicated. Peaks were identified with MACS2 (FDR<0.05). As expected, H3K4me3 is primarily found at promoters, H3K9me3 at intergenic regions (likely repeats) and H3K36me3 at gene bodies. The total number of peaks for each mark is indicated on the right. (**g**) PCA analysis of CUT&Tag read distribution in undifferentiated XX_ΔXic_ cells (circles) together with native ChIP-seq data(Żylicz et al., 2019) (triangles), previously generated in the parental TX1072 cell line in the same culture conditions (2iL). (**h**) Heatmap showing enrichment of CUT&Tag and ATAC-seq signals in chromatin states identified using *ChromHMM*. States were ordered according to the assigned identity shown on the right. (**i-j**) DNA accessibility and histone modifications in female XX_ΔXic_ mESCs prior to (Day 0) and at day 2 (i) and day 4 (j) of differentiation profiled by ATAC-seq and CUT&Tag. The tracks are overlaid in a way that an increased signal at day 0 and day 2/4 is colored in pink and orange es indicated, while the remaining signal is colored in black. Reads from two biological replicates were merged. Vertical bars below the tracks mark peaks identified in at least one time point and are colored in pink and orange, if the signal is significantly different (FDR<0.05) between the time points across both biological replicates. The screen results (Fig. 1) are shown below the tracks, where candidate REs that inhibit (blue) or activate (red) *Xist* expression in the negative or high fractions of the CRISPR screen are colored.

**Supplementary Figure 3.**
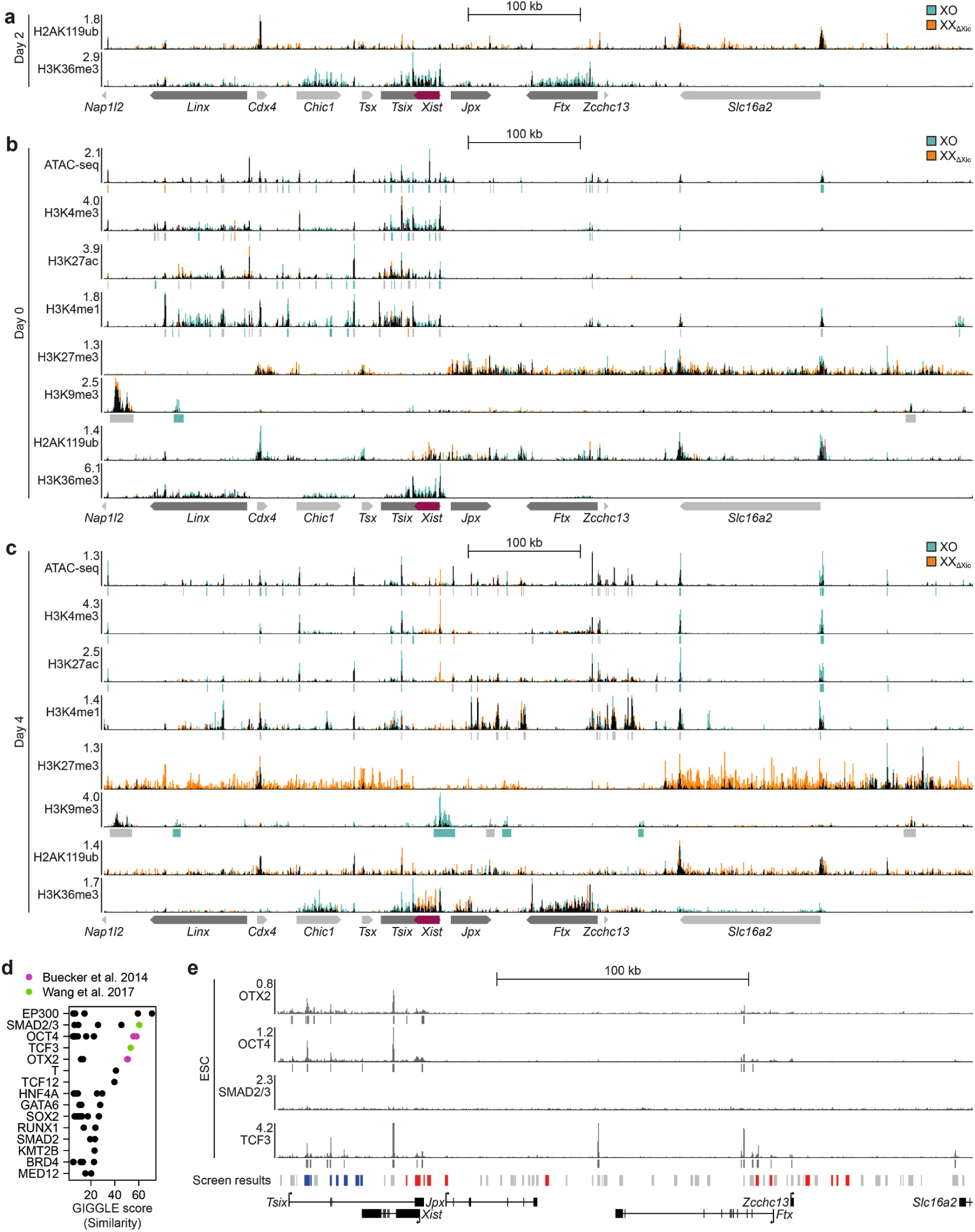
X-dosage information is decoded by promoter-proximal elements. (**a-c**) DNA accessibility and histone modifications in XX_ΔXic_ and XO mESCs prior to (b) and after 2 (a) or 4 days (c) of differentiation profiled by ATAC-seq and CUT&Tag. The tracks are overlaid in a way that an increased signal in XX_ΔXic_ and XO cells is colored in orange and teal as indicated, while the remaining signal is colored in black. Reads from two biological replicates were merged. Vertical bars below the tracks mark peaks identified in at least one time point and are colored in orange and teal, if the signal is significantly different (FDR<0.05) between cell lines across both biological replicates. (**d**) Enrichment of ChIP-seq signals in RE93,95,96,97 computed through the *Cistrome DB Toolkit*. The colored datasets are shown in (e) and in main Fig. 3e. (**e**) Published ChIP-seq profiles for OTX2, OCT4, SMAD2/3 and TCF3 in mESCs around the *Xist* locus (Buecker et al., 2014; Wang et al., 2017); positions where the signal extends beyond the depicted range are marked in red.

**Supplementary Figure 4.**
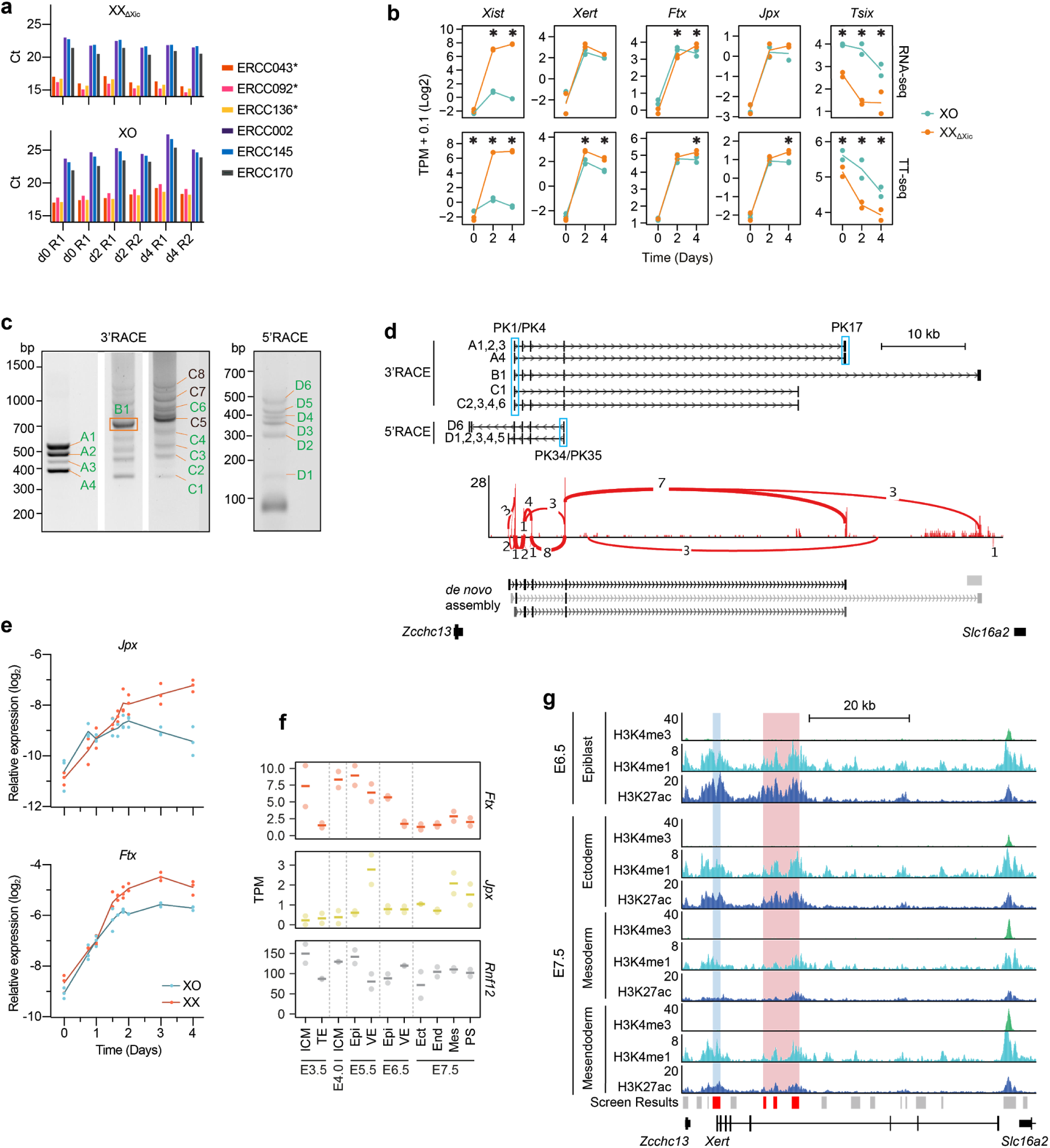
An unannotated enhancer-associated transcript is upregulated concomitantly with *Xist* at the onset of XCI. (**a**) Quality controls for TT-seq data showing enrichment of S4U-labelled spike-in RNAs (asterisks) compared to unlabelled controls after biotin-pulldown, assessed by RT-qPCR. (**b**) Quantification of nascent (bottom) and total RNA (top) of the indicated genes. Differential gene expression between XX_ΔXic_ and XO cells was assessed with DEseq (FDR<0.05, asterisks). (**c-d**) Isoform detection of *Xert*. (c) Agarose gel image of 3’- and 5’-RACE. Bands that were purified and Sanger-sequenced are labelled. Bands labelled in green gave successful Sanger sequencing results, which are summarized in (d, top). Sashimi plot and *de novo* transcript assembly (d, bottom) from polyA-RNA-seq data derived from differentiated (day 2) TX1072 cells. Numerical labels of the red lines indicate the number of split reads supporting the indicated splice junction. (**e**) Relative RNA expression of *Jpx* and *Ftx* in TX1072 XX and XO cells during differentiation, measured by RT-qPCR (n=3). (**f**) *Jpx, Ftx and Rnf12* RNA expression during early mouse embryonic development (E3.5-E7.5) from embryos of both sexes combined(Zhang et al., 2018). Inner cell mass (ICM), trophectoderm (TE), epiblast (Epi), visceral endoderm (VE), ectoderm (Ect), endoderm (End), mesoderm (Mes), primitive streak (PS). (**g**) ChIP-seq coverage tracks for H3K4me3, H3K4me1 and H3K27ac from E6.5 and E7.5 combined male and female mouse embryos at the *Xert* locus, derived from published datasets (Yang et al., 2019). Marked are the promoter/TSS region (blue) and enhancer cluster (pink). Horizontal bars in (f) and lines in (b,e) denote the mean of 2 (b,f) or 3 (e) biological replicates, dots represent individual measurements.

**Supplementary Figure 5.**
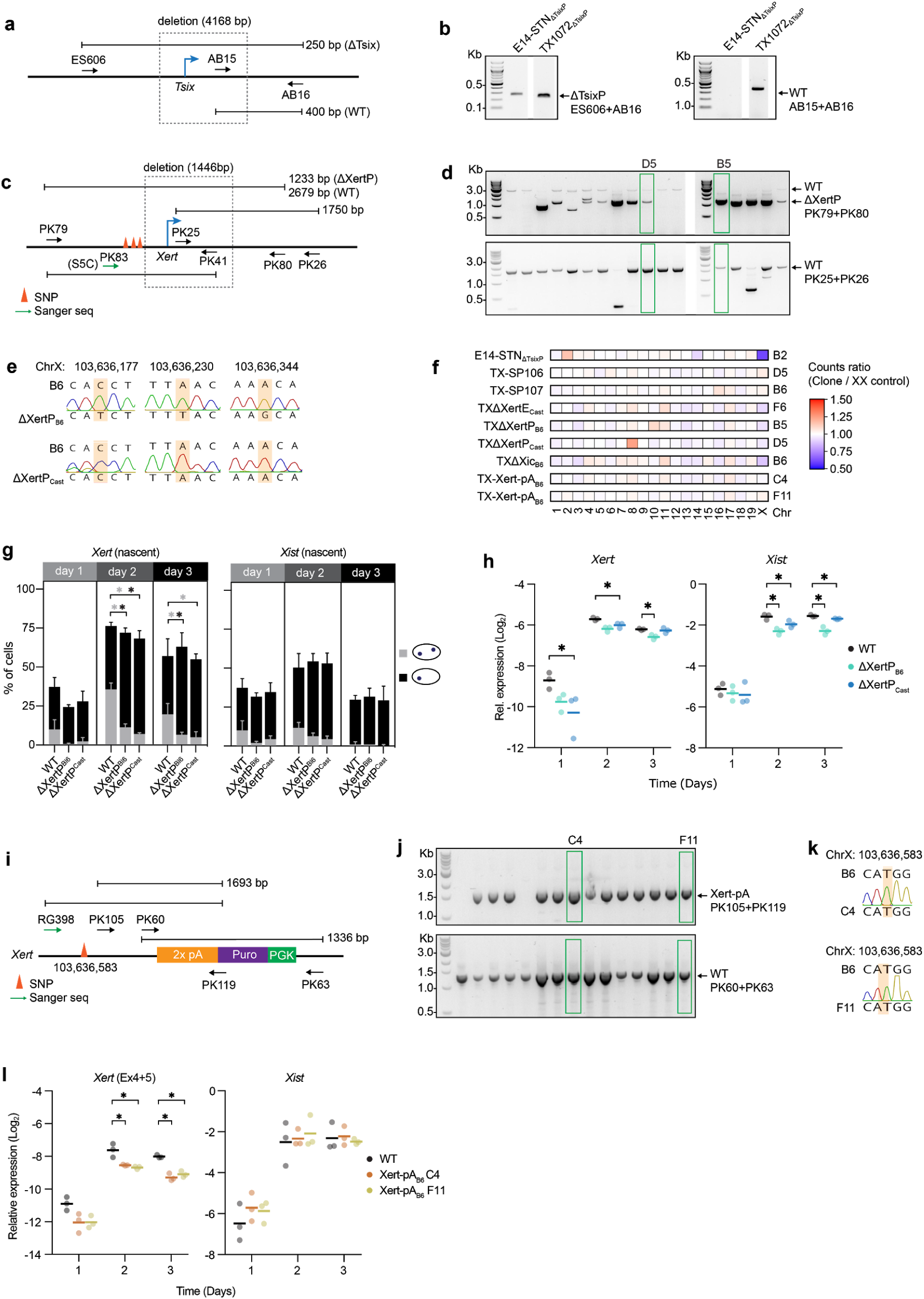
*Xert* transcription enhances *Xist* expression in *cis.* (**a-b**) Genotyping for E14-STN_ΔTsixP_ clones. (a) Arrows indicate primer positions. Lines denote PCR amplicons. (b) Agarose gel images of two genotyping PCRs (from E14-STN_ΔTsixP_ clones), that show the deleted (left panel) or the wt band only (right panel). (**c-e**) Genotyping of TXΔXertP clones. (c) Arrows indicate primer positions. Orange pyramids indicate the SNPs used in Sanger sequencing validation of the deleted allele. Lines indicate PCR amplicons. (d) Agarose gel images of two genotyping PCRs, that show the deletion and wt band (top) or the wt band only (bottom). Green boxes mark the clones that were selected for all further analyses. (e) Assessment of three SNPs (orange boxes) by sanger sequencing of amplicon PK79/PK41 and PK83 as sequencing primer (see c). Chromatogram and the genomic coordinates (mm10) of the SNPs are shown for both heterozygous ΔXertP lines. (**f**) Heatmap of NGS karyotyping data for all cell lines used in the study. Counts mapping to each chromosome were normalized to an XX control cell line. (**g-h**) Quantification of *Xert* and *Xist* RNA expression by RNA-FISH (g) and RT-qPCR in two heterozygous ΔXertP lines and parental TX1072 control line (WT). (**i-k**) Genotyping of Xert-pA insertion lines. (i) Arrows indicate primers positions, orange pyramids indicate the SNPs used for identification of the inserted allele by Sanger sequencing. Lines represent PCR amplicons. (j) Agarose gel images of two genotyping PCRs (from Xert-pA insertion clones), that show the insertion band (top) or the wt band (bottom). Green box marks the clones selected for further experiments. (k) Assessment of a SNP at the indicated genomic position (mm10) (orange box) by Sanger sequencing of amplicon RG398/PK119 and RG398 as sequencing primer (see i) for two selected Xert-pA clones. (**l**) *Xert* and *Xist* expression in two heterozygous Xert-pA clones (C4 and F11) and the parental TX1072 control (WT) assessed by RT-qPCR (l). For qPCR a position downstream of the insertion site was assayed for *Xert* (intron spanning from exon 4 and 5). In (h,l) horizontal bars denote the mean of 3 biological replicates, dots represent individual measurements. In (g) mean and s.d. of 3 biological replicates are shown. Asterisks indicate significance of *p*<0.05 using an unpaired two-tailed T-test.

**Supplementary Figure 6.**
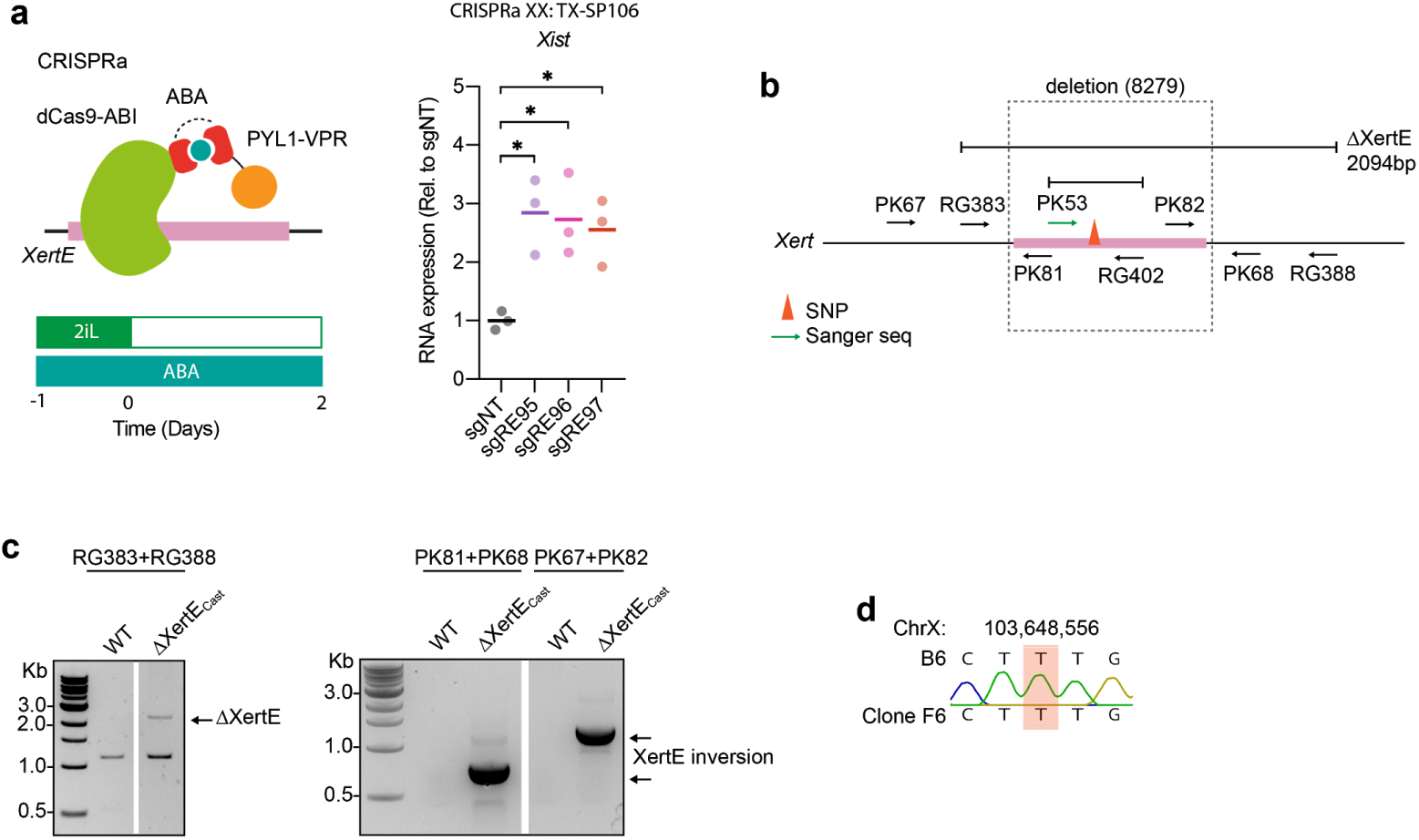
An intronic enhancer cluster within *Xert* activates *Xist* expression in *cis*. (**a**) Ectopic XertE activation in TX-SP106 cells stably expressing an inducible CRISPRa system and three sgRNA from a multiguide expression vector that target one RE or non-targeting controls (NT) (left). Relative *Xist* RNA expression after two days of differentiation was assessed by RT-qPCR (n=3) and normalized to sgNT, where each sgNT replicate is given by the geometric mean of four different sgNT plasmids. (**b-d**) Genotyping of ΔXertE mESC line, carrying a deletion of the *Xert* enhancer cluster on one allele and an inversion on the other. (b) Arrows indicate primer positions. The orange pyramid indicates the SNP used to identify the wt allele by Sanger sequencing in (d). Lines indicate PCR amplicons. (c) Agarose gel images of two genotyping PCRs of the ΔXertE clone used for further analyses, showing the deletion band (left) as well as an inversion on the other allele (right). (d) Sanger sequencing of amplicon PK53/RG402 and PK53 as sequencing primer (see b) to identify the deleted allele in ΔXertE mESCs. Chromatograms of the assessed SNP (orange box) are shown with its genomic coordinate (mm10).

**Supplementary Figure 7.**
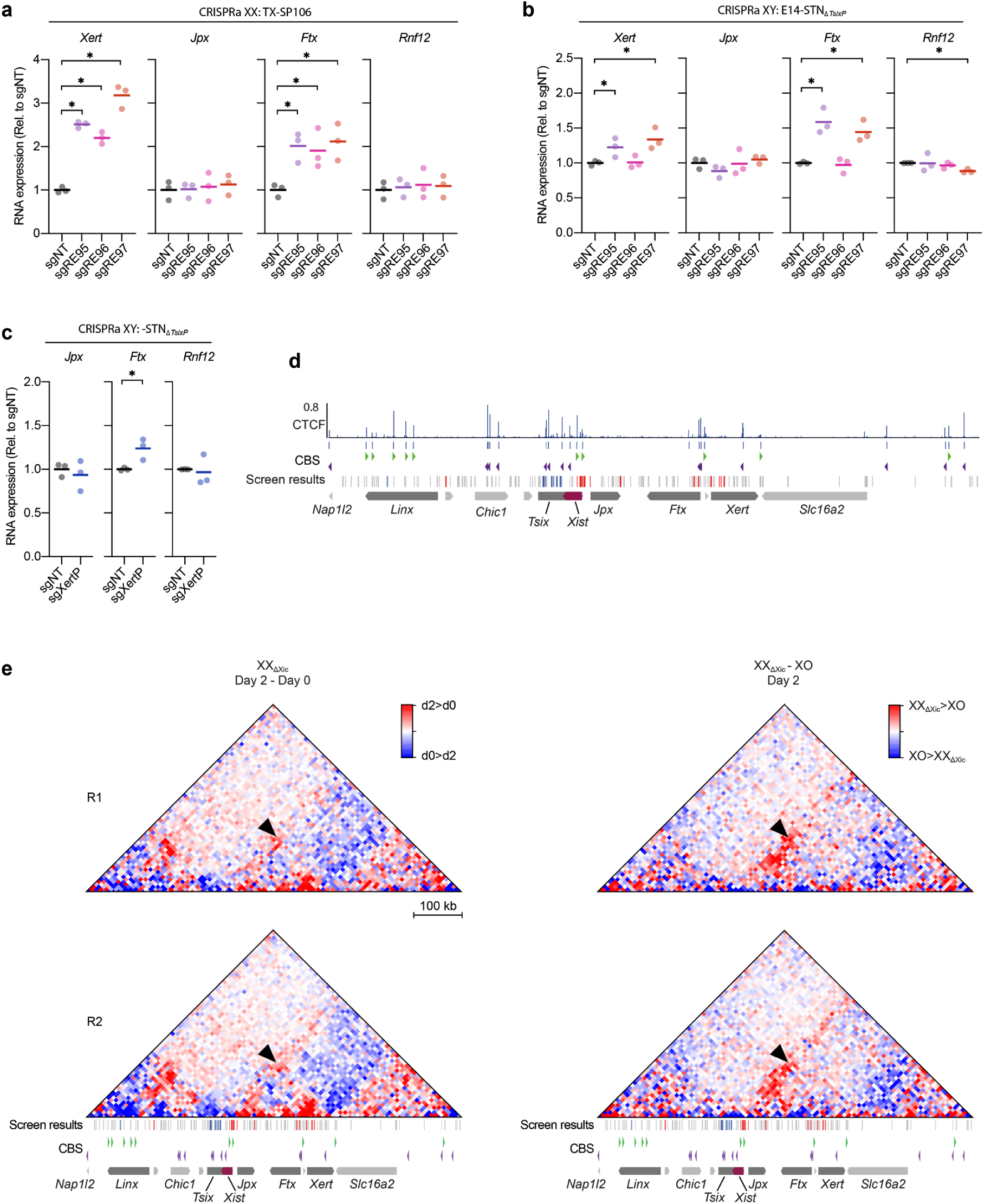
*Xert* and *Ftx* form a regulatory hub that exhibits increased contacts with the *Xist* promoter during initiation of XCI. (**a-c**) Ectopic activation of XertE (a-b) and XertP (c) through inducible CRISPR activation, using an ABA-inducible dCas9-VPR system (TX-SP106) in female (b) or a dox-inducible SunTag system in male (c) mESCs (E14-STN_ΔTsixP_). RNA expression was assessed in cells stably expressing three sgRNAs from a multiguide expression plasmid or non-targeting controls (NT) after 2 days of differentiation by RT-qPCR (n=3) and normalized to sgNT, where each sgNT replicate is given by the geometric mean of four different sgNT plasmids. Horizontal bars denote the mean of 3 biological replicates, dots represent individual measurements. Asterisks indicate significance of *p*<0.05 using an unpaired two-tailed *t*-test. (**d**) CTCF coverage track in male mESCs, derived from a published dataset(Stadler et al., 2011). Significant CTCF binding sites (CBS) are shown below the track and motif orientation is shown as triangles. The screen results are shown below the tracks, where candidate REs that inhibit (blue) or activate (red) *Xist* expression in the negative or high fractions of the CRISPR screen are colored. (**e**) Subtraction heatmaps as in Fig. 7g for two individual biological replicates (R1, R2) comparing day 0 and 2 in XX_ΔXic_ cells (left) and XX_ΔXic_ and XO cells at day 2 (right).

### Supplementary Tables

**Supplementary Table 1**. ATAC-seq/STARR-seq alignment and CRISPRi screen library design. Related to Fig. 1 and Supplementary Fig. 1.

**Supplementary Table 2**. CRISPRi screen library analysis. Related to Fig. 1 and Supplementary Fig.1.

**Supplementary Table 3**. CUT&Tag alignment and analysis. Related to Fig. 2, 3 and Supplementary Fig. 2, 3.

**Supplementary Table 4**. TT-seq,RNA-seq and Capture Hi-C alignment and analysis. Related to Fig. 4, 7 and Supplementary Fig. 4, 7.

**Supplementary Table 5**. *Xert* genomic regions and ORFs

**Supplementary Table 6**. Cell lines, oligos, probes, antibodies used in this study

### Supplementary screen discussion

#### Comparison of CRISPR screen results with published mutant phenotypes

While the *cis*-regulatory landscape of *Xist* has been dissected with genetic tools for more than 30 years (Galupa and Heard, 2018), we have now applied, for the first time, a high-throughput approach to allow comprehensive RE identification and quantification throughout the entire *Xic*. The CRISPRi screen we have performed relies on inactivation of promoter or enhancer elements through ectopic deposition of H3K9me3. As opposed to a genomic deletion, some RE classes cannot be targeted by CRISPRi, such as CTCF-bound insulators (Tarjan et al., 2019). Moreover, genomic resolution could be limited by the fact that H3K9me3 can spread over several kilobases around the targeted site (Thakore et al., 2015). Nevertheless our screen results are largely in agreement with previously reported mutant phenotypes.

REs previously reported to repress *Xist* were generally enriched in the Xist-high and depleted in the Xist-negative population, including the major *Tsix* promoter (RE49) (Lee et al., 1999), its enhancers *DxPas34* (RE50) (Cohen et al., 2007) and *Xite* (RE45-RE47) (Ogawa and Lee, 2003) and the LinxE element (RE12) (Galupa et al., 2020). Moreover, the screen identified the promoter regions of known *Xist* activators *Jpx* (RE61) (Tian et al., 2010), *Ftx* (RE85-88) (Chureau et al., 2011; Furlan et al., 2018) and *Rnf12* (RE127) (Barakat et al., 2011; Jonkers et al., 2009). For Ftx only very small effects were observed for the major TSS (RE88), but a strong phenotype was associated with RE85, which lies 6.5 kb downstream of the major TSS and acts as promoter of a minor isoform (Furlan et al., 2018). Importantly, RE85 and RE88 lie within the previously reported *Ftx* promoter deletions and in the region that has previously been targeted by CRISPRi (Furlan et al., 2018; Soma et al., 2014). Only one previously reported *Xist* RE, namely LinxP (RE20) (Galupa et al., 2020) was not identified within our screen, maybe due to a CRISPRi-insensitive mechanism of action. Moreover, two elements exhibited a strong phenotype in the screen, but no observable effect has been reported upon deletion, namely RE59 upstream of the *Xist* TSS and the major *Tsix* promoter (RE49) (Cohen et al., 2007; Newall et al., 2001). Since both regions lie in close proximity (<1.5kb) to very strong REs (Dxpas34, XistP), the screen phenotype might be attributed to H3K9me3 spreading in those cases. The pluripotency factor binding site in *Xist* intron 1 (RE55) (Barakat et al., 2011; Minkovsky et al., 2013; Navarro et al., 2008) exhibited an unusual pattern with being (weakly) depleted from both the negative and the high fractions, suggesting a dual function, where it inhibits initial Xist upregulation, but enhances expression levels, once Xist starts to be expressed.

The promoter-proximal elements RE57 and RE58 include the core promoter and ∼2kb of exon 1. This region has previously been implicated in *Xist* regulation and contains the somatic promoter P2, which is embedded in a CpG island (Johnston et al., 1998; Norris et al., 1994). Part of this region also encodes the repeat A of the *Xist* RNA, which is crucial for *Xist*’s gene silencing function (Wutz et al., 2002). Genomic deletion of the repeat A has been shown to abolish *Xist* upregulation both *in vitro* and *in vivo* (Hoki et al., 2009; Royce-Tolland et al., 2010). It is however difficult to distinguish, whether this phenotype is associated with a regulatory DNA element or with the inability of the *Xist* RNA to silence its *cis*-repressor *Tsix* (Mutzel et al., 2019; Robert-Finestra et al., 2020). Another part of the promoter-proximal region is bound by CTCF, YY1 and REX1, with the *Xist* activator YY1 and the repressor REX1 competing for overlapping binding sites (Chapman et al., 2014; Makhlouf et al., 2014; Navarro et al., 2006). Deletion of YY1 binding sites has been shown to abolish *Xist* upregulation (Makhlouf et al., 2014), also supporting the importance of RE57 in *Xist* regulation. Taken together, the screen is in excellent agreement with the literature on regions that regulate Xist expression within the *Xic*.

